# Formation of giant ER sheets by pentadecanoic acid causes lipotoxicity in fission yeast

**DOI:** 10.1101/2024.08.15.608187

**Authors:** Yojiro Hoshikawa, Natsuho Shirota, Hiroshi Tsugawa, Satoshi Kimura, Akihisa Matsuyama, Yoko Yashiroda, Hideaki Kakeya, Makoto Arita, Reiko Iizumi, Minoru Yoshida, Shinichi Nishimura

## Abstract

Excess amounts of saturated fatty acids are toxic to organisms, a condition termed lipotoxicity, which is often accompanied by pleiotropic cellular and tissue dysfunction. Here we show that pentadecanoic acid (C15:0) exerts toxicity on the fission yeast *Schizosaccharomyces pombe* by generating an aberrantly planar endoplasmic reticulum (ER) structure, which we named a “giant ER sheet.” Untargeted lipidomics revealed that C15:0 is incorporated into complex lipids depending on an acyl-CoA ligase Lcf1 and an acyl-CoA transferase Slc1, thereby increasing the saturation level of the acyl chains. The toxicity and giant ER sheet formation were abolished by deleting Lcf1 or Slc1, indicating that the incorporation of C15:0 into glycerophospholipids causes giant ER sheet formation. The giant ER sheets disrupted the correct migration of Mid1, a protein determining the cell division site, and physically blocked septum formation, hindering correct cell separation. Our results suggest that the ER is the primary site targeted by saturated fatty acids, leading to lipotoxicity.

## Introduction

Fatty acids (FAs) are fundamental molecules that constitute organisms and are included in diet. They function as energy storage molecules and signaling molecules, are used in post-translational modifications of proteins, and are components of membrane lipids. Despite their essentiality for living organisms, excess FAs cause sickness through cell and tissue dysfunction and cell death, a phenomenon known as lipotoxicity ^1–3^. The amount and species of FAs are crucial; factors such as chain length, degree of unsaturation, position and geometry of the double bonds, and oxidation status play significant roles ^4^. To investigate the function of these complex fatty acids, chemical genetics using lipid-modulating small molecules is expected to be a suitable approach.

The ER is an organelle which functions in diverse processes, including protein synthesis and transport, lipid metabolism, and calcium storage ^5, 6^. The ER forms an intricate network composed of at least two distinct membrane domains: sheets and tubules, each associated with specific functions. The sheets, which have flat structures with low curvature, are enriched with ribosomes on their cytoplasmic surface, indicating that protein synthesis occurs mainly in this region ^7, 8^. The tubules have high curvature and are presumed to be the major site for lipid synthesis and calcium regulation. Many proteins involved in the ER morphogenesis have been identified, and some are known to be related to diseases, although their molecular functions largely remain to be clarified ^6^. Furthermore, the contribution of the membrane lipids to the ER morphogenesis is not well understood, despite the fact that the majority of the lipid metabolism occurs at the ER ^9^.

While exploring natural products that perturb the cellular membrane function, we discovered that pentadecanoic acid (C15:0) potently inhibits growth of the fission yeast *Schizosaccharomyces pombe*, likely through dysregulating the membrane function. The mode-of-action of C15:0 was investigated using a combination of cytological analyses, untargeted lipidomics, and genetics. C15:0 was found to exhibit toxicity by inducing a giant ER sheet, an abnormally large ER structure, whose formation was abolished by depleting specific lipid-metabolic enzymes. The untargeted lipidomics implied that the giant ER sheet is induced by an increased saturation level of the acyl chains in glycerophospholipids (GPLs). Moreover, we demonstrate that myristic acid (C14:0) also induces the giant ER sheet and exhibits toxicity in certain lipid metabolism mutants. We propose that the giant ER sheet is one of the characteristic features leading to the pleiotropic phenotypes of lipotoxicity, which might be conserved in eukaryotes.

## Results

### Screen for membrane-targeting molecules to find the growth inhibitory activity of pentadecanoic acid

Cells lacking ergosterol biosynthetic genes are often tolerant to antimicrobial agents that target membrane lipids, not only ergosterol but also other lipid species ^10–14^. To identify novel membrane-targeting compounds, we conducted a screen using a wild-type strain and an ergosterol biosynthetic mutant strain (*erg31*Δ *erg32*Δ; **Extended Data Fig. S1**) of fission yeast. We screened a collection of 4160 fractions prepared from culture extracts of 416 marine bacteria and identified an active fraction derived from the culture extract of *Micromonospora* sp. Fractionation by HPLC furnished an active substance, whose chemical structure was deduced to be 2,3-dihydroxypropyl 14-methylpentadecanoate by NMR and MS analyses (**Fig. 1a**). This monoacylglycerol was previously named aggreceride B, which was isolated from *Streptomyces* sp. as an inhibitor of platelet aggregation with an unknown molecular mechanism ^15^.

**Figure 1.**
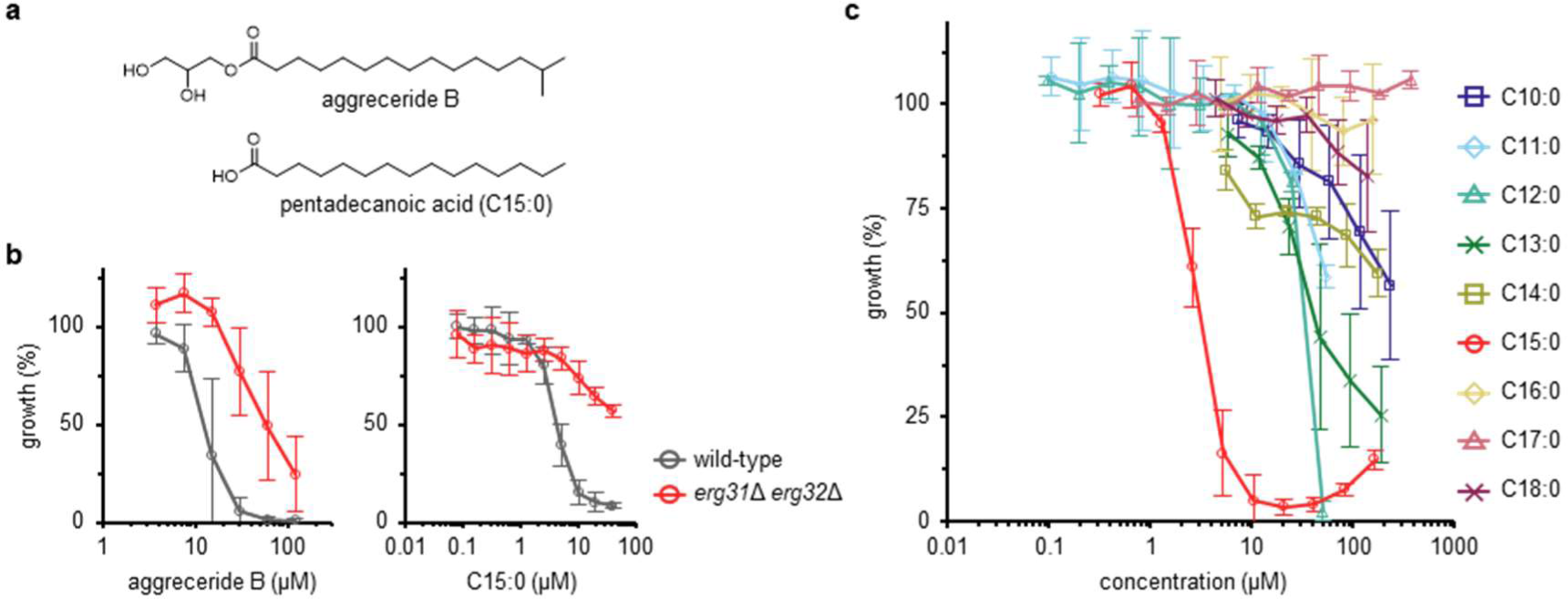
Aggreceride B and C15:0 inhibit the growth of S. pombe cells. **a,** Chemical structures of aggreceride B and C15:0. **b,** Aggreceride B and C15:0 sensitivity of wild-type and *erg31* Δ *erg32* Δ cells. n = 3. Error bars = SD. **c,** FA sensitivity of wild-type cells. Straight-chain saturated FAs with carbon numbers of 10 to 18 were tested. C10:0, C13:0, C14:0, C16:0, C18:0, n = 4; C11:0, C12:0, C15:0, C17:0, n = 3. Error bar = SD.

Aggreceride B showed potent growth inhibition against wild-type *S. pombe* cells, which was attenuated in *erg31*Δ *erg32*Δ cells (**Fig. 1b**). To determine the pharmacophore, we conducted a structure-activity relationship study of aggreceride B using a series of synthetic monoacylglycerols and commercially available FAs. Among monoacylglycerols tested, 2,3-dihydroxypropyl pentadecanoate showed potent growth inhibition comparable to that of aggreceride B, suggesting that the terminal methyl group of aggreceride B is not requisite (**Fig. S1a**). Furthermore, the glycerol moiety was not required for exhibiting activity, as pentadecanoic acid (C15:0; **Fig. 1a**) was found to be a potent growth inhibitor at almost the same level (**Fig. 1c**). Similar to aggreceride B, C15:0 exhibited weaker or no activity against mutants lacking ergosterol biosynthetic genes such as *erg2*, *erg31* and *erg32*, and *erg5* (**Fig. 1b, Extended Data Fig. S1b-c**). Mutants lacking *erg4* and *erg6* which are both responsible for the reaction at the branched methyl group in the side chain were susceptible to C15:0 (**Extended Data Fig. S1b-c**).

FAs other than C15:0 showed decreased or no activity (**Fig. 1c**): C14:0 exhibited only moderate activity, while palmitic acid (C16:0) was inactive, suggesting that the chain length of FAs is strictly recognized by cells. C15:0 showed no growth inhibition against other yeast species, including the fission yeasts *Schizosaccharomyces japonicus* and *Schizosaccharomyces octosporus*, and the budding yeast *Saccharomyces cerevisiae* (**Extended Data Fig. S2**). Taken together, C15:0 was shown to inhibit the growth of the fission yeast *S. pombe* in a highly specific manner, likely by disturbing the membrane function.

### Pleiotropic cytological defects induced by C15:0

To elucidate the mode of action of C15:0, we conducted cytological analyses. *S. pombe* cells are rod-shaped and undergo cytokinesis followed by septation at the center of the cell. We found that C15:0 induces two types of abnormal septa: *mid1* Δ-like septa and curved septa (**Fig. 2a,b**). Mid1 is one of the proteins that constitute the contractile actomyosin ring (CAR), and its deletion leads to misplacement of the division site and/or formation of multiple septa ^16, 17^. The curved septum is a novel phenotype, to our knowledge (see below). The population of cells with septa increased from 20% to 37% after three hours of C15:0 treatment, with 60% of the septa being abnormal (**Fig. 2c,d**). The percentage of cells with septa increased in a time-dependent manner: 78% after 6 hours of treatment, of which 69% had abnormal septa (**Fig. 2c,d**).

**Figure 2.**
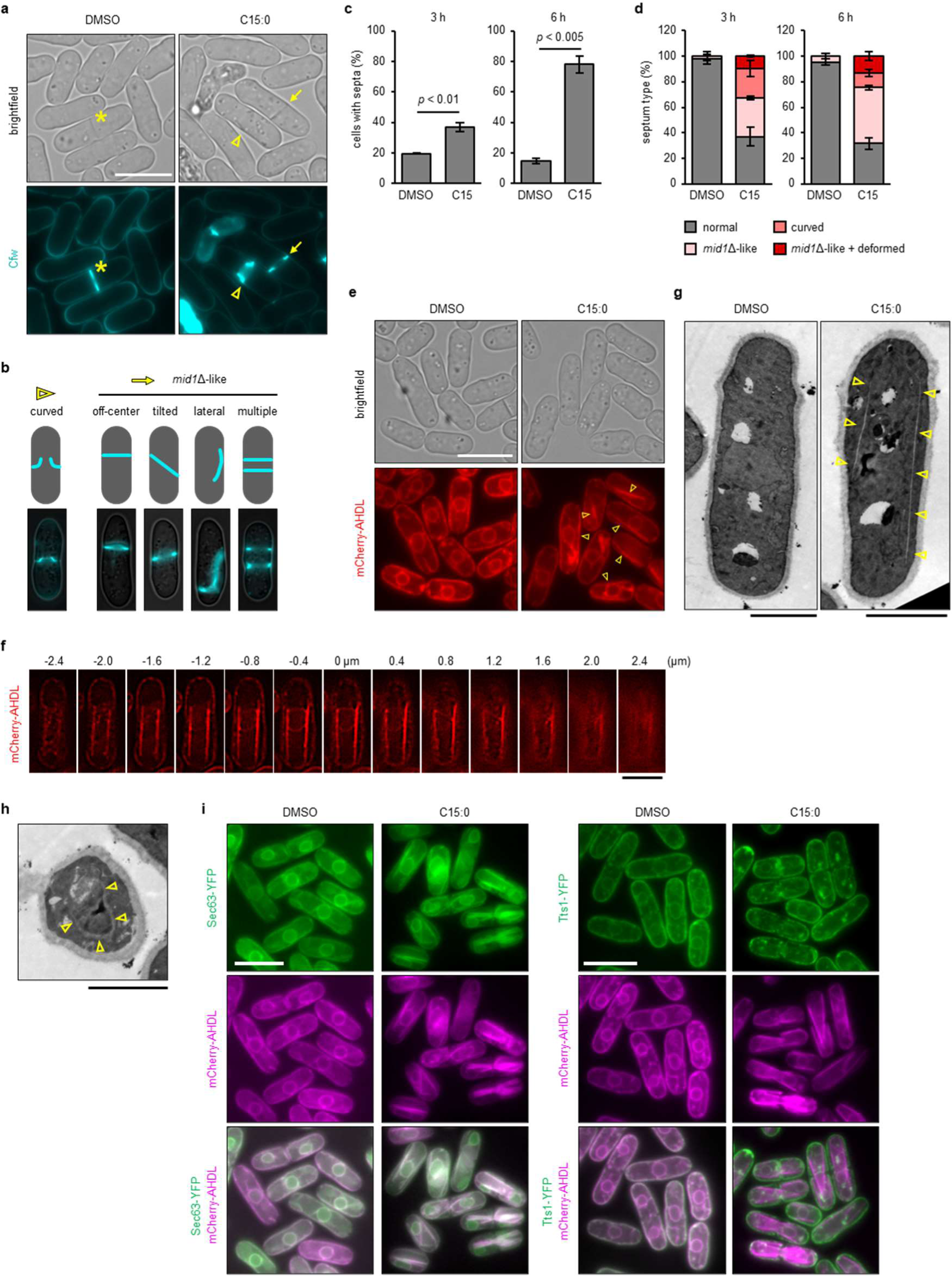
C15:0 induces various cytological defects. **a,** Visualization of septa with calcofluor white (Cfw). The asterisk, the arrowhead and the arrow indicate normal septum, curved septum and *mid1* Δ-like septum, respectively. Scale bar = 10 µm. **b,** Schematic and representative images of curved and *mid1* Δ-like septa. **c,d,** Fractions of cells with septum (**c**) and septum type (**d**). Cells were treated with DMSO (control) or C15:0 for three and six hours. n > 200 cells (**c**) and n >40 septa (**d**) from three independent experiments. Error bar = SD. P-values determined by unpaired two-tailed t-test. **e,** The ER visualized with mCherry-AHDL. Arrowheads indicate abnormal ER structures. Scale bar = 10 µm. **f,** Sequential z-stack images (0.2 µm gap) of the abnormal ER structure induced by C15:0. Background signal was removed using ImageJ. Scale bar = 5 µm. **g,** TEM image of control and C15:0 treated cells. Arrowheads indicate abnormal ER structures. Scale bar = 2 µm. **h,** TEM cross-section of a C15:0 treated cell perpendicular to the long axis of the cell. Yellow arrowheads indicate the abnormal ER structure. **i,** Localization of YFP-tagged Sec63 (a sheet localizing protein) and Tts1 (a tubule-forming protein). mCherry-AHDL was coexpressed to visualize the ER. Scale bar = 10 µm. Sec63-YFP and Tts1-YFP were expressed from pDUAL constructs ^47, 50^. Scale bar = 10 µm.

Mid1 localizes to the nucleus during interphase and migrates to the cell cortex near the nucleus upon mitotic entry ^18, 19^. We investigated the effect of C15:0 on the cellular distribution of Mid1-GFP in early mitotic cells in which mCherry-tagged α-tubulin (mCherry-Atb2) is polymerized in the nuclei. Dotted signals of Mid1-GFP were concentrated at the medial region under the control condition. In contrast, the distribution was dispersed by C15:0 treatment (**Extended Data Fig. S3**), which may be responsible for the *mid1* Δ-like septa.

During the microscopic analyses using Mid1-GFP, we noticed the presence of aberrantly shaped nuclei in C15:0 treated cells (**Extended Data Fig. S4a**). The nuclear deformation was confirmed with a nuclear envelope (NE) marker, Cut11-mCherry (**Extended Data Fig. S4b**). These results prompted us to investigate whether C15:0 affects the progression of mitosis. The dynamics of the spindle pole body (SPB) marked by Sid4-GFP were observed in synchronized *cdc25-22* cells. C15:0 treatment did not affect the speed of SPB segregation, but a delay in the onset of SPB segregation was detected (**Extended Data Fig. S4c,d**). Notably, 45% of cells failed to properly distribute nuclei after cell division, likely due to a combination of delayed mitosis and imperfect cytokinesis (**Extended Data Fig. S4e,f**). The mis-segregation of nuclei should be one of the causes of the growth inhibition by C15:0.

The NE is continuous with the ER. We next analyzed the morphology of the ER by expressing an ER luminal marker, mCherry-AHDL ^20^. The fluorescent signal was detected around the nuclei and the cell cortex under the control condition, as reported previously ^20^. In contrast, C15:0-treated cells exhibited unnaturally linear signals along the long axis of the cells, observed in nearly 80% of the cell population (**Fig. 2e**). Although the structure resembled microtubules, mCherry-Atb2 did not overlap with the abnormal ER structure (**Extended Data Fig. S5a**). Instead, the linear ER signals were observed continuously in multiple focal planes (**Fig. 2f**), indicating its planar architecture. The planar structures in C15:0-treated cells were also observed by transmission electron microscopy (TEM) (**Fig. 2g**). We found that the electron density of the structure was low, suggesting that the protein concentration of the lumen is low. Notably, round structures were observed in the slice perpendicular to the long axis of the cells, confirming the sheet structure of the abnormal ER (**Fig. 2h**). Thus, we refer this abnormal ER structure as a giant ER sheet.

The giant ER sheet colocalized with YFP-tagged Sec63, a protein presumed to reside mainly at the sheet domain of the ER with low membrane curvature (**Fig. 2i**)^21^. In contrast, ER tubule-forming proteins Yop1 and Tts1 were excluded from the giant ER sheet (**Fig. 2i, Extended Data Fig. S5b**) ^20, 22–24^. These results support the idea that the abnormal ER generated by C15:0 has a sheet structure.

Since the ER and the plasma membrane (PM) are physically tethered to regulate the lipid homeostasis of the membranes ^25^, we visualized PM sterols to investigate the effect of C15:0. Sterols are concentrated at cell tips or division sites in fission yeast cells ^26^. Line scan analyses revealed the attenuation of the polarized distribution of sterols in C15:0-treated cells (**Extended Data Fig. S6**). Taken together, C15:0 was shown to exhibit pleiotropic effects on cell morphology.

### The giant ER sheet causes growth inhibition

Since multiple morphological changes were observed, we decided to investigate their relationships to identify the phenotype relevant to the toxicity of C15:0. C15:0 generated curved septa, as described above (**Fig. 2a,b**). We visualized the septa and the ER simultaneously and found that the giant ER sheets were caught by the curved septa (**Extended Data Fig. S7a**). Time-lapse analyses revealed that the septa could not cleave the giant sheets but instead curved and grew along the giant sheets (**Fig. 3a**). In the TEM analyses, a rod-like structure, which could be derived from the giant ER sheets, was found to be placed between the growing septa (**Extended Data Fig. S7b**). We also tested the relationship between the giant ER sheets and the dispersion of the Mid1 distribution. When cells co-expressing Mid1-GFP and mCherry-AHDL were treated with C15:0, clusters of Mid1-GFP signals were observed exclusively with signals from the giant ER sheets (**Fig 3b**). Thus, the giant ER sheet is likely the upstream event of the abnormal septa, physically blocking the growth of the septa and disturbing the correct migration of Mid1 to the cell cortex.

**Figure 3.**
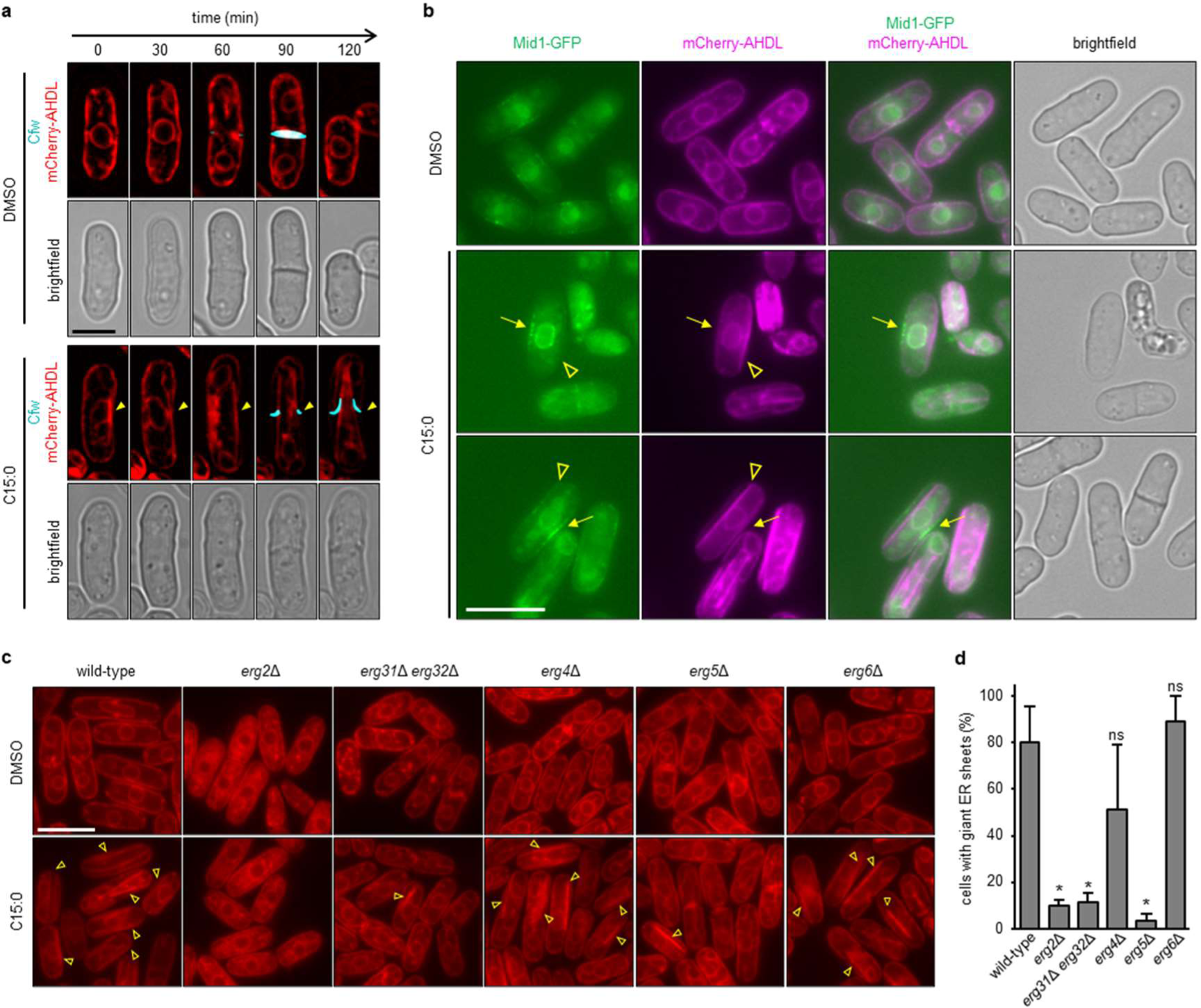
The generation of the giant ER sheet is the primary cause of the growth inhibition. **a,** Timelapse observation of the septum and the ER. The cells were observed every 30 minutes after two hours of treatment with DMSO or C15:0. The images are representative of each condition. The background signal was removed using ImageJ. Scale bar = 5 µm. **b,** Observation of the ER (mCherry-AHDL) and Mid1-GFP. Arrows indicate Mid1-GFP signal on the side opposite of the giant ER sheet (arrow heads). Scale bar = 10 µm. **c,** The ER in *erg* mutants. Scale bar = 10 µm. **d,** The rate of cells with the giant ER sheets. n > 300 cells from three independent experiments. Error bar = SD. P-values determined by unpaired two-tailed t-test. * p < 0.05.

Given that the ER and PM are tethered through the ER-PM contact sites, we wondered whether the giant ER sheet affects the polarity of sterols at the PM, probably through dysfunction of the contact site. To test this, we performed filipin staining using the *scs2*Δ *scs22*Δ cells, in which the ER-PM contact sites are defective and the cortical ER is largely detached from the PM ^27^. Interestingly, the polarity of the PM sterols disappeared in the *scs2*Δ *scs22*Δ strain regardless of C15:0 treatment (**Extended Data Fig. 8a,b**). This indicated that the ER-PM contact sites are crucial for the polarized distribution of the PM sterols, while the giant ER sheet likely induces dysfunction of the ER-PM contact sites. Notably, the giant ER sheet was still induced by C15:0 in the *scs2*Δ *scs22*Δ strain (**Extended Data Fig. S8c**), indicating that the ER-PM contact site and the cortical ER are not requisite for the induction of giant ER sheets.

Some ergosterol biosynthesis mutants are resistant to C15:0 (**Fig. 1B, Extended Data Fig. S1c**). We tested whether C15:0 can generate the giant ER sheets in these mutants (**Fig. 3c,d**). C15:0-sensitive *erg4*Δ and *erg6*Δ cells formed giant sheets upon C15:0 treatment, while C15:0-resistant *erg2*Δ, *erg31*Δ *erg32* Δ, and *erg5*Δ cells did not. There was a clear correlation between giant ER sheet formation and the toxicity of C15:0. Furthermore, the giant sheets were not formed in the C15:0-tolerant *S. cerevisiae* cells (**Extended Data Fig. S9**). Collectively, giant ER sheet formation seems to be the primary phenotype responsible for the growth inhibition by C15:0.

### C15:0 perturbs the yeast lipidome

As the ER is the major site of the lipid biosynthesis, we next examined whether C15:0 is incorporated into the membrane lipids at the ER, potentially altering the physical properties of the membrane to induce the giant ER sheet. To investigate this idea, we conducted LC-MS/MS-based untargeted lipidomics. Wild-type and *erg31* Δ *erg32* Δ cells were cultured with or without C15:0 for three or six hours, and then the total lipids were extracted from the harvested cells. Lipid samples were subjected to liquid chromatography coupled with quadrupole time-of-flight mass spectrometry (LC/Q-TOF-MS). The MS data were analyzed by MS-DIAL ^28^, to quantify 590 lipid species across subclasses (e.g. free FAs, phosphatidylcholine (PC), phosphatidylinositol (PI), sterol ester, etc.) (**Extended Data Table S1**).

We first performed hierarchical clustering and heatmap analyses to visualize the global changes in the lipidome (**Fig. 4**). Eight experimental conditions were divided into two clusters, DMSO treatment and C15:0 treatment. As expected, C15:0 caused significant changes in the yeast lipidome. Each cluster was further divided into two subclusters, wild-type and *erg31*Δ *erg32* Δ cells, indicating that the ergosterol biosynthesis has a broad impact on the lipidome. The molecular species were divided into seven clusters (**Extended Data Table S2**). Clusters I and VI were mainly composed of molecular species that were increased by C15:0 treatment. While the classes of the increased lipid species were diverse, they shared a common feature: the incorporation of C15:0 and odd-chain FAs (OCFAs) that could be derived from C15:0 (57 among 61 for cluster I and 92 among 93 for cluster VI; **Extended Data Table S2**). It was shown that *erg31* Δ *erg32*Δ cells can also uptake and incorporate C15:0. The increase of the incorporation of OCFAs smaller than C13:0 was rarely observed, possibly due to the lack of the β-oxidation pathway in *S. pombe* ^29^. Clusters II, III, IV, V, and VII were formed in a quantity-dependent manner (**Fig. 4**).

**Figure 4.**
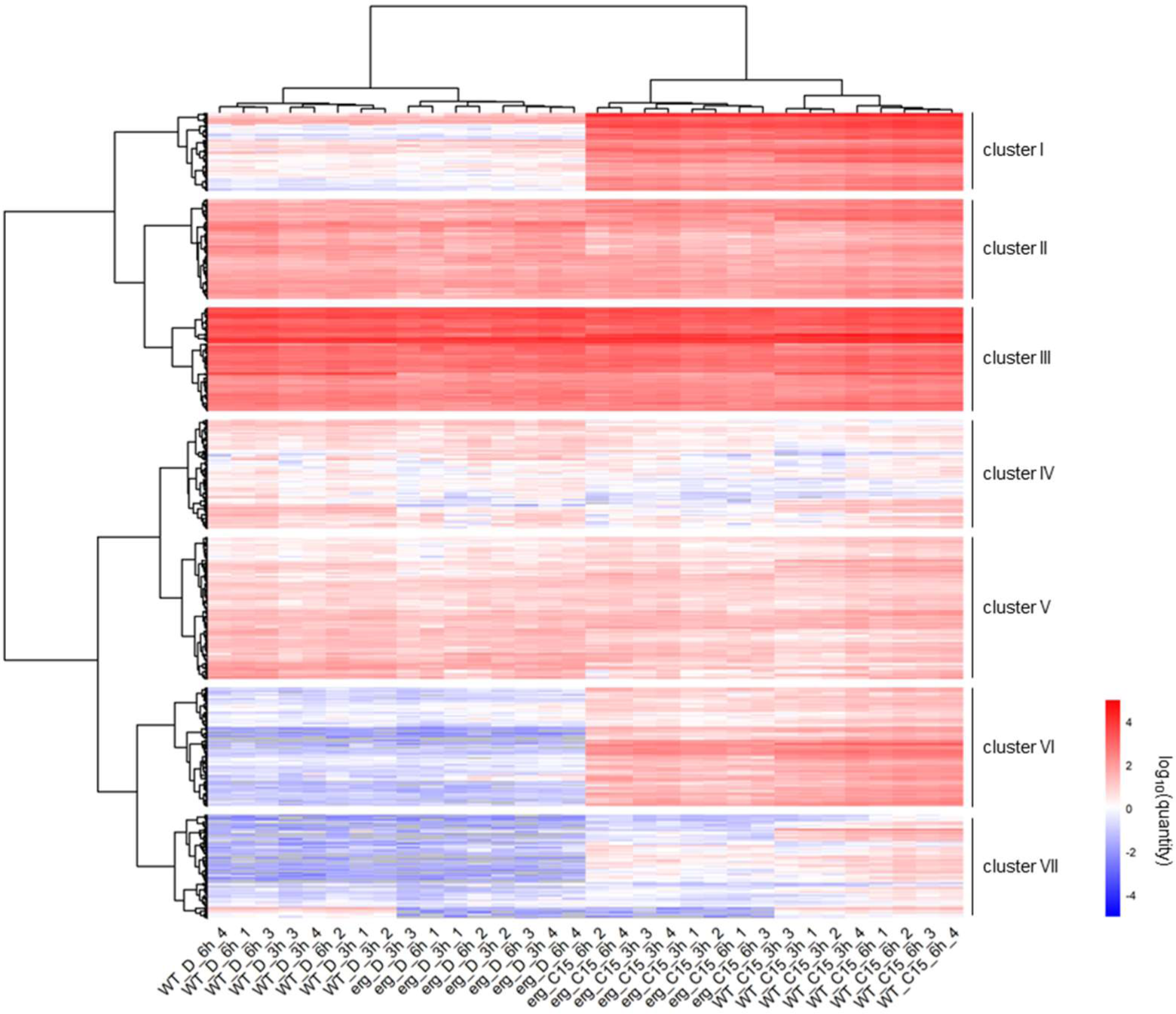
C15:0 causes global changes in yeast lipidome. The rows represent the lipid molecular species, and the columns represent the samples. The colors represent the quantity (pmol / 10^8^ cells) on a log_­_scale. Each sample name indicates the strain (wild-type, WT or erg31Δ erg32Δ, erg), the compound treated (DMSO, D or C15:0, C15), the time of the compound treatment (3 or 6 hours), and the sample number (1-4). The lipid molecular species are divided into seven clusters I - VII. See **Extended Data Table S2** for details.

When we carefully analyzed the MS/MS data, we noticed that cluster I contained molecular species with two C15:0 such as PC 15:0_15:0 and PI 15:0_15:0. Their retention times in the LC and the *m/z* values were same as PC 12:0_18:0 and PI 12:0_18:0, respectively, which were relatively abundant in DMSO-treated cells. Although the two overlapping molecular species could not be quantified separately, the MS/MS spectra unambiguously showed that two sets of C15:0 are incorporated in PC and PI in the C15:0 treated cells (**Extended Data Fig. S10**).

### Effect of C15:0 on the profile of free fatty acids and glycerophospholipids

To understand how lipidomes are altered by C15:0, we next investigated changes in molecular species within each lipid class. We first analyzed the profile of the free FAs (**Fig. 5a, Extended Data Fig. S11**). In wild-type cells, C18:1 (oleic acid), C18:0, and C16:0 were the three most abundant species, while C15:0 and C18:1 became the most abundant two after the C15:0 treatment. The amount of oleic acid increased concomitantly, suggesting that the oleic acid synthesis is up-regulated in response to the C15:0 treatment. Notably, the oleic acid level in *erg31* Δ *erg32* Δ mutant cells was already higher than that in wild-type cells, and their abundance of oleic acid prevented C15:0 from becoming the most abundant FA after the C15:0 treatment.

**Figure 5.**
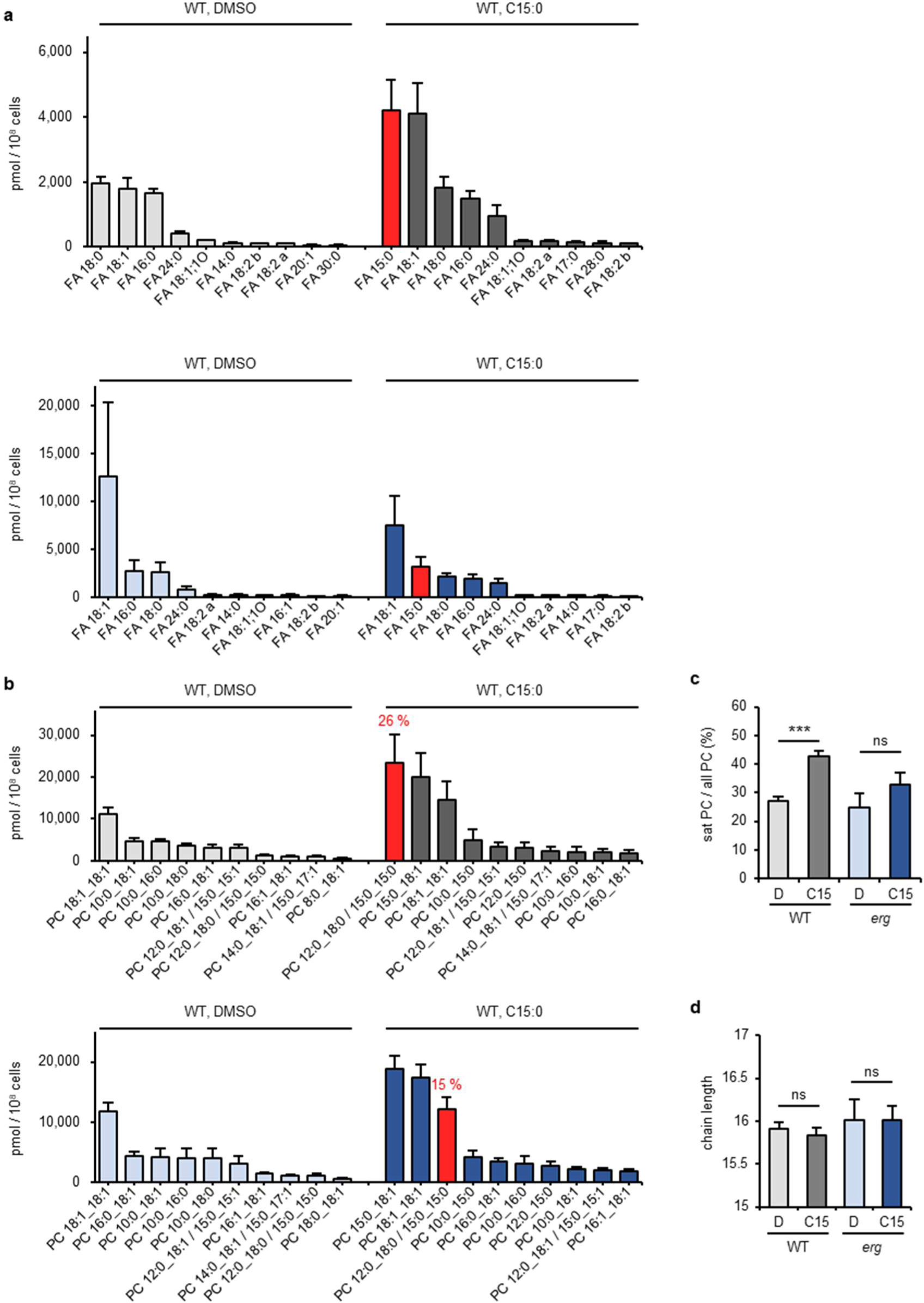
C15:0 perturbs the profile of free fatty acids and GPLs. **a,** Profile of free FAs at three-hour chemical treatment. The quantity (pmol/10^8^ cells) of top ten molecular species under indicated conditions are shown. **b,** Profile of PC at three-hour chemical treatment. The quantity (pmol/10^8^ cells) of top ten molecular species under indicated conditions are shown. The red bar indicates the species possessing two C15:0 as acyl chains and its proportion (%) is shown. PC 12:0_18:0 and PC 15:0_15:0 are shown together due to the limitations at LC separation. **c,** The proportion (%) of saturated PC among total PC at three-hour chemical treatment. **d,** The average carbon chain length of acyl chains in PC at three-hour chemical treatment. (**a-d**) *erg* means *erg31* Δ *erg32*Δ mutant. (**a-d**) n = 4. Error bar = SD。 (**c-d**) P-values determined by unpaired two-tailed t-test. ns, no significance; * p < 0.05; ** p < 0.01; *** p < 0.001.

Next, we focused on the acyl groups of GPLs. Among GPLs, PC is the major lipid class in the eukaryotic cell membrane, with PC 18:1_18:1 being the most abundant species in wild-type *S. pombe* cells. The C15:0 treatment made PC 15:0_15:0 the most abundant, followed by PC 15:0_18:1 (**Fig. 5b, Extended Data Fig. S12a**). PC 15:0_15:0 constituted nearly a quarter of the total PC, and over half of the saturated PC, which has two saturated acyl groups (**Fig. 5b, Extended Data Fig. S12a,b**). The rate of the saturated PC drastically increased, e.g. from 27% to 43% after three hours of C15:0 treatment (**Fig 5c, Extended Data Fig. S12c**). Notably, GPL species with two saturated long-chain acyls were rare either in wild-type and *erg31*Δ *erg32* Δ cells, while the acyl chains making up saturated GPLs consisted of one medium-chain FA (C6-12) and one long-chain FA (C13-C21). Changes in the average acyl chain length were subtle (**Fig. 5d**). These trends were alleviated in *erg31*Δ *erg32* Δ cells, although C15:0 was incorporated into PC species. For example, the level of PC 15:0_15:0 was smaller in *erg31* Δ *erg32* Δ cells than in wild-type cells (i.e. 25.8 ± 0.9% in wild-type and 15.4 ± 0.5% in *erg31*Δ *erg32* Δ cells, under three-hour treatment conditions) (**Fig. 5b, Extended Data Fig. S12a**). Additionally, there was no significant increase in saturated PC level by C15:0 treatment in *erg31* Δ *erg32*Δ cells (**Fig. 5c, Extended Data Fig. S12c**).

PI and phosphatidylserine (PS) profiles were similar to that of PC. In wild-type cells, C15:0 treatment significantly increased PI and PS with C15:0 as an acyl group, especially PI 15:0_15:0 and PS 15:0_15:0, and increased the proportion of saturated species (**Fig. S13a-c, S14a-c**). These changes were suppressed in *erg31*Δ *erg32*Δ cells. In contrast to PC, we observed a decrease in the average length of acyl chains of PI and PS (**Extended Data Fig. S13d, 14d**). This was particularly noticeable for PS, likely due to the absence of medium-chain FAs in the PS acyl chains (**Extended Data Fig. S14d**).

The phosphatidylethanolamine (PE) profile showed unique features, different from those of PC, PI, and PS. In wild-type cells, the majority of PE had two unsaturated acyl chains, while only 0.6% of PE comprised saturated PE species. In contrast, 20-30% of PC and PI species were saturated ones (**Extended Data Fig. S15a-c**). When wild-type cells were treated with C15:0, PE 15:0_18:1 was abundantly detected, while PE 15:0_15:0 was low (4.1 ± 0.4% of total PE) (**Extended Data Fig. S15a**). These results suggest a cellular system that maintains a relatively high level of unsaturated PE. In the *erg31*Δ *erg32* Δ cells, PE 15:0_15:0 formation was further reduced (1.3 ± 0.1% after three-hour treatment). The chain length of PE was reduced by C15:0 treatment, though the effect was small, likely due to the low incorporation of C15:0 into PE (**Extended Data Fig. S15d**). Taken together, the incorporation of C15:0 into GPLs such as PC, PI, and PS and the resulting increase in saturated GPLs correlate with the growth inhibition by C15:0, which can possibly be mitigated by supplementation with oleic acid.

### Acyl-CoA synthetase Lcf1 is necessary for the activity of C15:0

To investigate if the incorporation of C15:0 into GPLs is the cause of the toxicity, we examined the involvement of long-chain fatty acyl-CoA synthetases (FACSs). FACS is necessary for the cellular uptake of extracellular FAs and the metabolic activation of FAs ^30, 31^. *S. pombe* has two FACS homologs, Lcf1 and Lcf2. Simultaneous deletion of *lcf1* and *lcf2* causes a severe decrease in long-chain fatty acyl-CoA synthetase activity and a rapid loss of viability in the stationary phase, while single deletion is viable ^32, 33^. We tested two single-gene-deletion mutants for their sensitivity to C15:0 and found that *lcf1*Δ cells, but not *lcf2*Δ cells, are tolerant to C15:0 (**Fig. 6a**). The giant ER sheets were not generated in *lcf1*Δ cells but were in *lcf2*Δ cells upon treatment with C15:0 (**Fig. 6b, Extended Data Fig. S16a**). These results indicated that C15:0 is incorporated into GPLs after conversion to acyl-CoA by Lcf1, which is required for giant ER sheet formation and cell growth inhibition.

**Figure 6.**
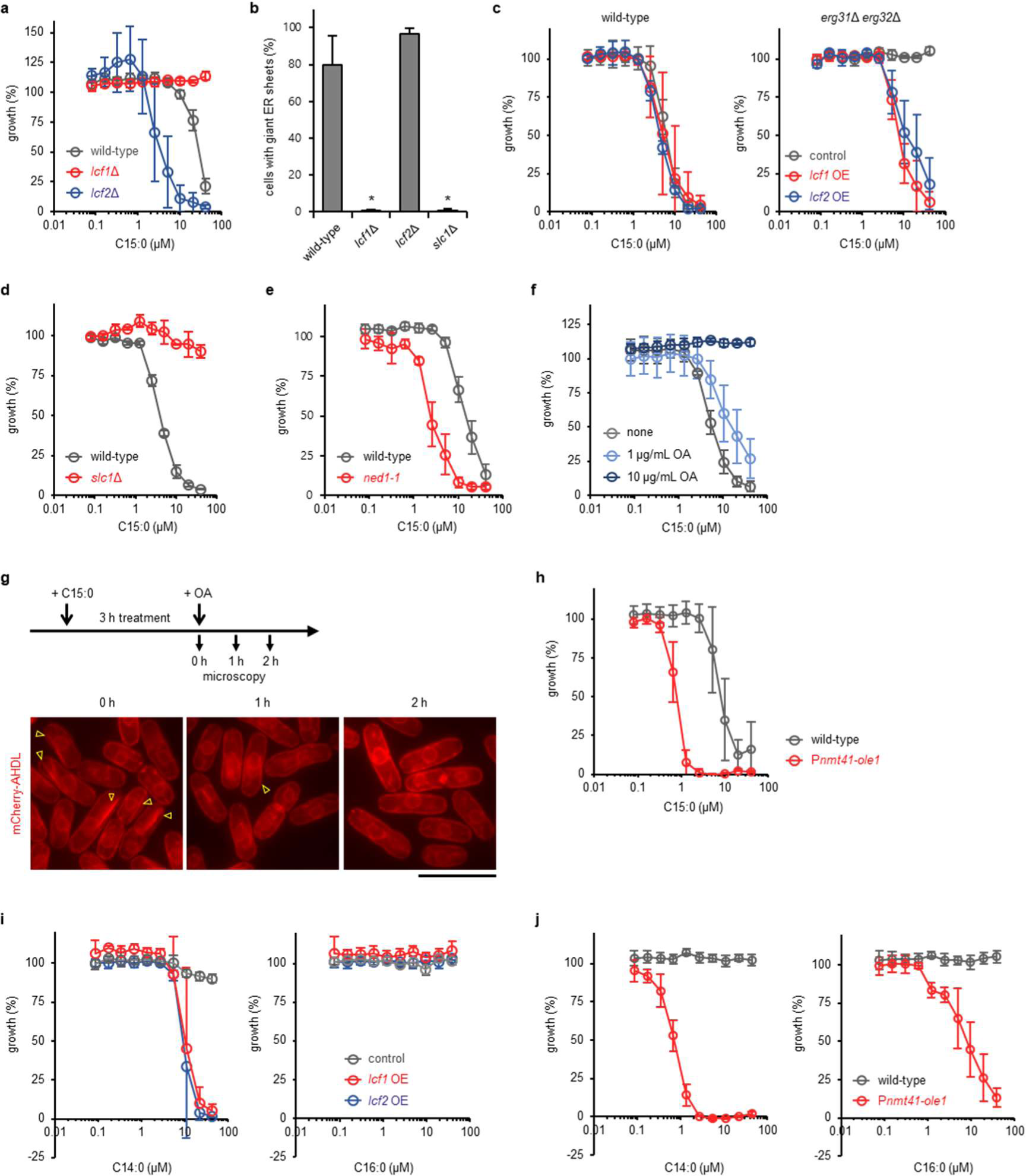
Involvement of Lcf1 and Slc1 in the action of saturated long-chain FAs. **a,** C15:0 sensitivity of *lcf1*Δ and *lcf2*Δ cells. Cells were cultured at 30°C. n = 3. Error bar = SD. **b,** The rate of cells with the giant ER sheets. n > 300 cells from three independent experiments. Error bar = SD. P-values determined by unpaired two-tailed t-test. ns, no significance; * p < 0.05. **c,** C15:0 sensitivity of wild-type and *erg31*Δ *erg32*Δ cells overexpressing *lcf1* or *lcf2* from multicopy plasmid pREP41. Control means empty vector, OE is overexpression. Cells were cultured in MM for 48 hours. n = 3. Error bar = SD. **d,** C15:0 sensitivity of *slc1*Δ cells. n = 3. Error bar = SD. **e,** C15:0 sensitivity of *ned1-1* cells. Cells were cultured at 30°C. n = 3. Error bar = SD. **f,** The activity of C15:0 in the culture supplemented with oleic acid. n = 3. Error bar = SD. **g,** Recovery of the morphological defect of ER by oleic acid. Cells were treated with C15:0 for three hours to induce the generation of the giant ER sheets before adding oleic acid to the culture. The observation was conducted every hour after oleic acid supplementation. Scale bar = 10 µm. **h,** C15:0 sensitivity of P*nmt41-ole1* cells. Cells were cultured in MM for 48 hours. n = 3. Error bar = SD. **i,** C14:0 and C16:0 sensitivity of cells overexpressing *lcf1-YFH* and *lcf2-YFH.* The strains from the ORFeome collection were used^47^. OE means overexpression. Cells were cultured in MM for 24 hours. n = 3. Error bar = SD. **j,** C14:0 and C16:0 sensitivity of P*nmt41-ole1* cells. Cells were cultured in MM for 48 hours. n = 3. Error bar = SD.

We next examined the effect of overexpression of FACSs. While the overexpression did not affect the C15:0-sensitivity in wild-type cells, the C15:0-resistance of *erg31*Δ *erg32* Δ was canceled by overexpressing *lcf1* or *lcf2* (**Fig. 6c**). Intriguingly, there was no difference between *lcf1* and *lcf2* overexpression in the degree of alteration of the C15:0 sensitivity. These results indicate that not only Lcf1 but also Lcf2 can recognize C15:0 as a substrate when overexpressed, although their physiological functions may differ in wild-type cells, since deletion of *lcf2*, but not *lcf1*, failed to confer tolerance to C15:0 (**Fig. 6a**). Additionally, these results suggest that the lower incorporation of C15:0 in the *erg31* Δ *erg32* Δ lipidome compared to wild-type lipidome is responsible for the lower sensitivity to C15:0 in the ergosterol mutant.

### Involvement of a putative *O*-acyl transferase Slc1 in the action of C15:0

We next conducted a focused screen employing mutant cells lacking *O* -acyltransferases, which function to incorporate acyl chains into complex lipid species. In the fission yeast genome, 18 genes encode *O* -acyltransferases: 17 are annotated with the gene ontology “*O* - acyltransferase activity (GO:0008374)” and one (*gpc1)* with “glycero-3-phosphocholine acyltransferase activity (GO:0106158)”. Five of them are predicted to encode acetyltransferases, which are unlikely to be functional for the acyl chain transfer to complex lipid species. Since three of the remaining 13 genes are essential, we examined the other ten genes using gene deletion mutants (**Extended Data Table S3**). Among the mutants tested, only one mutant showed altered sensitivity: the cell growth of *slc1*Δ was almost completely resistant to C15:0 (**Fig. 6d, Extended Data Fig. S16b**). Slc1 is a putative 1-acylglycerol-3-phosphate *O* -acyltransferase that converts lysophosphatidic acid (LPA) to phosphatidic acid (PA). The giant ER sheets were undetectable in C15:0-treated *slc1*Δ cells (**Fig 6b, Extended Data Fig. S16c**). It is noted that Ale1 and Vps66 are predicted to catalyze the same enzymatic reaction as Slc1, but the mutants lacking *ale1* or *vps66* gene did not show altered sensitivity to C15:0 (**Extended Data Fig. S16b**). This observation suggests a complex regulation system of lipid metabolism by three functional homologs.

The screen included *are1* and *are2*, encoding acyl-CoA-sterol acyltransferases required for sterol ester biosynthesis, and *dga1* and *plh1*, encoding acyltransferases required for triacylglycerol synthesis ^34^. Since Are1 and Are2, and Dga1 and Plh1 are functionally redundant, we tested double deletion mutants. In both cases, the minimal inhibitory concentrations of C15:0 were not changed (**Extended Data Fig. S16d-f**), ruling out the possibility that the incorporation of C15:0 into neutral lipids is the cause of cell growth inhibition. This scenario was supported by an experiment using a mutant of lipin, which converts PA into DG to generate neutral lipids. An increase in the membrane lipid synthesis was observed in the temperature-sensitive lipin mutant *ned1-1* ^35^. The *ned1-1* mutant showed higher sensitivity to C15:0 than wild-type cells (**Fig. 6e**). These results suggest that the cause of growth inhibition by C15:0 is the overproduction of membrane phospholipids that contain C15:0 as acyl groups.

### Oleic acid antagonizes the effect of C15:0

The lipidomics revealed that C15:0 increased the saturation level of the acyl chains in GPL, an effect that was attenuated in *erg31*Δ *erg32* Δ cells, which contain higher concentrations of oleic acid. To investigate the functional antagonism between C15:0 and oleic acid, we examined the effect of oleic acid on the sensitivity to C15:0. Growth inhibition by C15:0 was attenuated by 1 µg/ml of oleic acid and was completely canceled by 10 μg/ml of oleic acid (**Fig. 6f**). Additionally, the giant ER sheets disappeared with the supplementation of oleic acid (**Fig. 6g**).

We next manipulated the expression level of the *ole1* gene, which encodes the sole, putative Δ-9 FA desaturase in fission yeast, responsible for the production of oleic acid. By inserting the *nmt41* promoter ^36^ upstream of the *ole1* gene, we successfully decreased the mRNA level of *ole1* (**Extended Data Fig. S16g**). As expected, P*nmt41-ole1* mutant showed increased sensitivity to C15:0 (**Fig. 6h**). Taken together, we conclude that the increase in the saturation level of GPLs is the cause of giant ER sheet formation and growth inhibition by C15:0.

### The giant ER sheet can be induced by C14:0 in sensitized mutants

Finally, we tested whether the generation of the giant ER sheet was limited to C15:0-treated cells. Since FAs other than C15:0 were inactive or showed only moderate inhibitory activity against the wild-type strain, we used strains that showed increased sensitivity to C15:0 (**Fig. 6c,h**). As expected, overexpression of Lcf1-YFP or Lcf2-YFP sensitized cells to C14:0, comparable to the effect of C15:0 (**Fig. 6i**). Importantly, the giant ER sheets were generated by C14:0 in these cells (**Extended Data Fig. S17**). Additionally, P*nmt41-ole1* cells also became sensitive to C14:0 (**Fig. 6j**). The efficacy of C16:0 was smaller than those of C14:0 and C15:0 (**Fig. 6j**), possibly because C16:0 is the precursor of oleic acid and therefore detoxified by desaturation. Notably, the giant ER sheets were observed in P*nmt41* -*ole1* cells even in the absence of exogenous FAs (**Extended Data Fig. S18**). These results support the idea that the giant ER sheet is a significant factor in lipotoxicity induced by long-chain, saturated FAs.

## Discussion

In this study, we found that a FA C15:0 induces cell growth inhibition with unique phenotypes, such as abnormal septum formation, improper positioning of the division plane, depolarized distribution of sterols at the PM, and deformed nuclear morphology in the fission yeast *S. pombe.* We also found that C15:0 treatment induces characteristic rearrangement of the ER structure, which we named the giant ER sheet, physically blocking cell division. Our untargeted lipidomics indicates that C15:0 is actively incorporated into complex lipid species, drastically changing the lipidome and generating giant ER sheets. Giant ER sheet formation should be responsible for at least part of the C15:0-induced toxicity, as deleting genes for specific enzymes involved in the incorporation of FAs into complex lipids prevented giant ER sheet formation and rescued cell growth in the presence of C15:0. Our findings extend to C14:0: exogenous C14:0 induced giant ER sheet formation and inhibited cell growth in some lipid mutants.

Lipotoxicity can be observed in mammalian cells as well as in unicellular eukaryotes such as yeast ^37, 38^. Experiments using cultured mammalian cells have demonstrated that the saturated FA palmitic acid (C16:0) induces mitochondrial dysfunction and ER stress, leading to apoptosis ^37^. Angular-shaped dilated ER was observed in mammalian cells when excess amounts of C16:0 were added ^39^, although the consequences of this abnormal structure are not known. In the budding yeast *S. cerevisiae*, treatment with C16:0 under conditions that suppress oxidative reactions (i.e. heme synthase deficiency and hypoxia) reduced the degree of unsaturation of the acyl chains of complex lipids, resulting in ER expansion and polygonal ER/NE formation ^40, 41^. Recent studies have shown that polygonal ER/NE formation is toxic, as it is accompanied by nuclear rupture. Since giant ER sheet formation by excess saturated FAs appears to be conserved among eukaryotes, it is worth investigating the relationships between giant ER sheet formation and cellular dysfunction, to understand the molecular basis of lipotoxicity.

The fact that a single double bond significantly changes the physical properties of FA (e.g., melting temperature 69.6 °C for C18:0 and 13-16 °C for oleic acid) suggests that giant ER sheet formation is caused by the excessive incorporation of saturated FAs into complex lipids ^39–41^. In this study, genetic analysis indicated that the incorporation of C15:0 into GPL, not neutral lipids, is required for the toxicity of C15:0 (**Fig. 6f**). GPLs with unsaturated acyl chains are most concentrated in the ER among the organelles of the secretary pathway ^42^, making the ER highly susceptible to reduction in GPL unsaturation. Untargeted lipidomics revealed that the occupancy of saturated PE among all PE remained low under all conditions (**Extended Data Fig. S15**). Considering that the GPLs abundant in the ER are PC, PI, and PE ^43^, the increased saturation of PC and PI may contribute to the change in ER structure. This implies that the proper saturation level of PC and PI determines the shape of the ER and likely regulates biological events at the ER.

We showed that the toxicity and giant ER sheet formation by C15:0 is mediated by specific enzymes among multiple functional homologs in fission yeast. Only Lcf1, but not Lcf2, and Slc1 are specifically involved in susceptivity to C15:0. Lcf1 and Slc1 may have substrate specificity that prefers C15:0, or alternatively, C15:0 cannot be detoxified by Ole1, a desaturase. Ole1 might desaturate C14:0 and C16:0 directly or after their elongation to C18:0, rendering them non-toxic oleic acid. Both scenarios align with the finding that C14:0 and C16:0 induced growth inhibition under *ole1* knockdown conditions. These possibilities should be carefully examined in future studies.

Genes encoding enzymes responsible for downstream reactions in the ergosterol-biosynthesis pathway are nonessential for fission yeast growth: mutants with precursor sterols or their metabolic derivatives, instead of ergosterol are viable ^10, 44^. Some of these ergosterol-biosynthesis mutants were tolerant to C15:0 (**Fig. 1b, Extended Data Fig. S1c**). Untargeted lipidomics revealed an increased level of oleic acid in *erg31* Δ *erg32* Δ cells compared to wild-type cells (**Fig. 5a, Extended Data Fig. S11**), which likely competes with C15:0. Multiple phenotypes have been reported for ergosterol-biosynthesis mutants, previously interpreted as results of increased immature sterol species. However, some might be due to the increase in oleic acid. For example, we previously reported that *erg31*Δ *erg32* Δ cells are tolerant to antifungal natural products, heronamides, ^12^ which target phospholipids with saturated hydrocarbon chains. The higher level of oleic acid in the mutant could increase unsaturated GPLs, preventing heronamide binding to the membrane. Lipidomics revealed that endogenous C15:0 is virtually absent in *S. pombe*, and exogenously added C15:0 inadvertently hijacks the GPL biosynthesis pathway, leading to its incorporation into GPLs. By identifying the enzyme that misrecognizes C15:0, it may be possible to develop a new experimental system to actively incorporate saturated fatty acids into GPLs. Detailed analyses of the effects of giant ER sheet formation on ER function may elucidate the relationship between the intricate structure and function of the ER.

## Methods

### Yeast strains and cultivation

Yeast strains used in this study are listed in **Extended Data File**. *S. pombe* strains prepared in this study were generated either by genetic cross, by a PCR-based strategy, or by transformation with plasmids ^45, 46^. In the case of the PCR-based strategy, gene deletion mutants were made by replacing the coding region with kanMX6 module in pFA6a-kanMX6, and the insertion of P*nmt41* at the upstream region of *ole1* was carried out as described previously using a plasmid pFA6a-kanMX6-P41nmt1 ^46^. For mutants overexpressing *lcf1* or *lcf2* from extrachromosomal plasmid pREP41-lcf1 or pREP41-lcf2, leucine auxotroph cells were transformed with the plasmids and transformants were selected by growing on a medium lacking leucine. Otherwise, *lcf1* or *lcf2* was overexpressed at the *leu1* locus under the control of the *nmt1* promoter ^47^.

Cells were cultured in either YES medium composed of yeast extract, glucose, and 225 mg/L each of five supplements (Ade, Ura, L-Leu, L-His, and L-Lys) or Edinburgh minimal medium 2 (EMM) ^48^. For selection, we supplemented YES with 100 µg/ml G418 (500 µg/mL for EMM), 100 µg/ml nourseothricin, 100 µg/ml hygromycin B, and 15 µg/ml blasticidin-S. For suppressing the gene expression utilizing P*nmt41* promoter, we cultured cells in YES or EMM supplemented with 2 µM thiamine.

### Recombinant DNA and oligonucleotides

Recombinant DNA and oligonucleotides used in this study are listed in **Extended Data File.** pREP41-lcf1 and pREP41-lcf2 were constructed in this study. First, ORF of *lcf1* or *lcf2* was amplified by PCR using *S. pombe* genomic DNA as a template, and primers lcf1-pREPclon-F and lcf1-pREPclon-R or lcf2-pREPclon-F and lcf2-pREPclon-R, respectively. The obtained DNA fragments were mixed with *Nde* I digested pREP41 and the mixture was introduced into competent cells of the *E. coli* DH5α to clone *lcf1* or *lcf2* ORF into pREP41 by in vivo *E. coli* cloning (iVEC). The sequences of the plasmids were confirmed by DNA sequencing (Fasmac).

### Screening and purification of aggreceride B

As general experimental procedures, LC-MS analysis was carried out using an Agilent 6530 Accurate-Mass Q-TOF mass spectrometer coupled to an Agilent 1260 LC system. NMR spectra were recorded on a JEOL 500 MHz instrument. ^1^H and ^13^C chemical shifts are shown relative to the residual solvent: *δ*_­_7.26 and *δ*_­_77.0 for CDC_­_. Chemical shifts (*δ*) are shown in parts per million (ppm) and coupling constants (*J*) are in hertz (Hz). Chemicals for organic synthesis were purchased from Sigma and used without further purification.

For the screening, A collection of fractions (4,160), which were obtained by SiO_­_gel column chromatography of marine microbial culture extracts, was screened using an anti-yeast assay. In this assay, we used two fission yeast strains: a wild-type and a mutant strain lacking *erg31* and *erg32* genes. Among fractions tested, one fraction derived from *Micromonospora* sp. CNY045 showed a preferred selectivity in which the fraction exhibited growth inhibition against wild-type cells while showed no activity against the mutant cells. A portion of the hit fraction (0.48 mg) was subjected to RP-HPLC (Phenomenex, Luna 5u C18(2) 100A, 10 x 250) using a gradient elution system (2 ml/min of aq MeCN, 20% for 12 min, 20 to 100% in 78 min, 100% for 30 min.). A fraction with the activity of the purpose was further separated by RP-HPLC using another gradient system (2 ml/min of aq MeCN, 70% for 12 min, 70 to 100% in 78 min, 100% for 30 min.). One of the active fraction was analyzed by LC-MS and NMR to reveal that the active substance was aggreceride B: ^1^H NMR (CDCl_­_, 500 MHz) *δ* 4.21(dd, *J* = 11.5, 4.5 Hz, 1H), 4.15 (dd, *J* = 11.5, 6.5 Hz, 1H), 3.94 (m, 1H), 3.70 (dd, *J* = 11.5, 4.0 Hz, 1H), 3.60 (dd, J = 11.5, 6.0 Hz, 1H), 2.35 (t, *J* = 7.8 Hz, 2H), 1.63 (m, 2H), 1.51 (m, 1H), 1.29-1.25 (ovl, 18H), 1.15 (m, 2H), 0.86 (d, *J* = 7.0 Hz, 6H); HR ESI-MS *m/z* 353.2664 [M + Na]^+^ calcd for C_­_H_­_NaO_­_, 353.2662.

### Synthesis of monoacylglycerols

Taking MAG-C15 as an example, firstly, pentadecanoic acid (124.7 mg, 0.52 mmol), DMAP (13.8 mg, 0.11 mmol), and DCC (133.0 mg, 0.65 mmol) were added to a stirred solution of glycerol (200 uL, 2.7 mmol) in DCM (5 mL) and DMF (1 mL). After being stirred for 23 h, the reaction was stopped by the addition of H_­_O (20 mL). The mixture was extracted with EtOAc (20 mL, three times). The organic layers were combined, washed with brine, dried over Na_­_SO_­_, and concentrated. The mixture was subjected to a SiO_­_column (hexane/EtOAc = 4:1 and 1:1) to obtain a crude fraction containing the target compound. This fraction was passed through the ODS short column using MeOH as a mobile phase, and the obtained solution was concentrated (139.9 mg). A portion of this fraction (16 mg) was subjected to RP-HPLC (Phenomenex Luna 5 μm C18(2), φ10 x 250 mm) using a gradient elution system (70 to 100% aq MeCN, 2 ml/min) to obtain MAG-15 (13.0 mg, 70.0%) as a white solid; ^1^H NMR (CDCl_­_, 500 MHz) *δ* 4.18 (dd, *J* = 11.5, 4.5 Hz, 1H), 4.13 (dd, *J* = 11.5, 6.0 Hz, 1H), 3.92 (m, 1H), 3.67 (dd, *J* = 11.5, 3.0 Hz, 1H), 3.58 (dd, *J* = 11.5, 5.5 Hz, 1H), 2.93 (brs, 1H), 2.55 (brs, 1H), 2.34 (t, *J* = 7.5 Hz, 2H), 1.61 (m, 2H), 1.24-1.28 (ovl, 22H), 0.87 (t, *J* = 7.0 Hz, 3H); ^13^C NMR (CDCl_­_, 125 MHz) *δ* 174.4, 70.2, 65.1, 63.3, 34.1, 31.9, 29.6 (5C), 29.4, 29.3, 29.2, 29.1, 24.9, 22.7, 14.1; HR ESI-MS *m/z* 339.2506 [M + Na]^+^ calcd for C_­_H_­_NaO_­_, 339.2512. Other monoacylglycerols were synthesized similarly and their structures were confirmed by HRMS and NMR analysis.

### Physicochemical properties of synthesized monoacylglycerols

#### MAG-C10

A white solid; ^1^H NMR (CDCl_­_, 500 MHz) *δ* 4.17 (dd, *J* = 11.5, 5.0 Hz, 1H), 4.13 (dd, *J* = 12.0, 6.0 Hz, 1H), 3.92 (m, 1H), 3.68 (dd, *J* = 11.5, 3.5 Hz, 1H), 3.58 (dd, *J* = 11.5, 5.5 Hz, 1H), 2.34 (t, *J* = 7.5 Hz, 2H), 1.61 (m, 2H), 1.25-1.28 (ovl, 12H), 0.87 (t, *J* = 7.0 Hz, 3H); ^13^C NMR (CDCl_­_, 125 MHz) *δ* 174.4, 70.2, 65.1, 63.3, 34.1, 31.8, 29.4, 29.2 (2C), 29.1, 24.9, 22.6, 14.1; HR ESI-MS *m/z* 269.1719 [M + Na]^+^ calcd for C_­_H_­_NaO_­_, 269.1723.

#### MAG-C13

A white solid; ^1^H NMR (CDCl_­_, 500 MHz) *δ* 4.18 (dd, *J* = 11.5, 4.5 Hz, 1H), 4.13 (dd, *J* = 11.5, 6.0 Hz, 1H), 3.92 (m, 1H), 3.68 (dd, *J* = 11.5, 3.5 Hz, 1H), 3.58 (dd, *J* = 11.5, 5.5 Hz, 1H), 2.93 (brs, 1H), 2.56 (brs, 1H), 2.34 (t, *J* = 7.5 Hz, 2H), 1.61 (m, 2H), 1.25-1.28 (ovl, 18H), 0.87 (t, *J* = 7.0 Hz, 3H); ^13^C NMR (CDCl_­_, 125 MHz) *δ* 174.4, 70.2, 65.1, 63.3, 34.1, 31.9, 29.6 (3C), 29.4, 29.3, 29.2, 29.1, 24.9, 22.6, 14.1; HR ESI-MS m/z 311.2197 [M + Na]^+^ calcd for C_­_H_­_NaO_­_, 311.2193.

#### MAG-C14

A white solid; ^1^H NMR (CDCl_­_, 500 MHz) *δ* 4.20 (dd, *J* = 11.5, 4.5 Hz, 1H), 4.15 (dd, *J* = 11.5, 6.0 Hz, 1H), 3.93 (m, 1H), 3.69 (dd, *J* = 11.5, 4.0 Hz, 1H), 3.60 (dd, *J* = 11.5, 6.0 Hz, 1H), 2.35 (t, *J* = 7.5 Hz, 2H), 1.62 (m, 2H), 1.25-1.29 (ovl, 20H), 0.88 (t, *J* = 7.0 Hz, 3H); ^13^C NMR (CDCl_­_, 125 MHz) *δ* 174.4, 70.2, 65.1, 63.3, 34.1, 31.9, 29.7, 29.62 (2C), 29.57, 29.4, 29.3, 29.2, 29.1, 24.9, 22.7, 14.1; HR ESI-MS *m/z* 325.2351 [M + H]^+^ calcd for C_­_H_­_NaO_­_, 325.2349.

#### MAG-C16i

A white solid; ^1^H NMR (CDCl_­_, 500 MHz) *δ* 4.21 (dd, *J* = 11.5, 4.5 Hz, 1H), 4.15 (dd, *J* = 11.5, 6.9 Hz, 1H), 3.93 (m, 1H), 3.70 (dd, *J* = 11.5, 4.0 Hz, 1H), 3.60 (dd, *J* = 11.5, 6.0 Hz, 1H), 2.35 (t, *J* = 7.5 Hz, 2H), 1.63 (m, 2H), 1.51 (m, 1H), 1.29-1.25 (ovl, 18H), 1.14 (m, 2H), 0.86 (d, *J* = 6.5 Hz, 6H); ^13^C NMR (CDCl_­_, 125 MHz) *δ* 174.4, 70.3, 65.1, 63.3, 39.0, 34.1, 29.9, 29.71, 29.66, 29.63, 29.58, 29.4, 29.2, 29.1, 28.0, 27.4, 24.9, 22.7 (2C); HR ESI-MS *m/z* 353.2660 [M + H]^+^ calcd for C_­_H_­_NaO_­_, 353.2662.

#### MAG-C16

A white solid; ^1^H NMR (CDCl_­_, 500 MHz) *δ* 4.21 (dd, *J* = 12.0, 4.5 Hz, 1H), 4.15 (dd, *J* = 12.0, 6.5 Hz, 1H), 3.93 (m, 1H), 3.70 (dd, *J* = 11.5, 4.0 Hz, 1H), 3.60 (dd, *J* = 11.5, 5.5 Hz, 1H), 2.35 (t, *J* = 7.5 Hz, 2H), 1.63 (m, 2H), 1.32-1.25 (ovl, 24H), 0.88 (t, *J* = 7.0 Hz, 3H); ^13^C NMR (CDCl_­_, 125 MHz) *δ* 174.4, 70.3, 65.2, 63.3, 34.1, 31.9, 29.7-29.6 (5C), 29.58, 29.44, 29.35, 29.2, 29.1, 24.9, 22.7, 14.1; HR ESI-MS *m/z* 353.2654 [M + H]+ calcd for C_­_H_­_NaO_­_, 353.2662.

#### MAG-C18

A white solid; ^1^H NMR (CDCl_­_, 500 MHz) *δ* 4.21 (dd, *J* = 11.5, 4.5 Hz, 1H), 4.15 (dd, *J* = 11.5, 6.5 Hz, 1H), 3.93 (m, 1H), 3.70 (dd, *J* = 11.5, 4.0 Hz, 1H), 3.60 (dd, *J* = 11.5, 5.5 Hz, 1H), 2.35 (t, *J* = 7.5 Hz, 2H), 1.63 (m, 2H), 1.29-1.25 (ovl, 28H), 0.88 (t, *J* = 7.0 Hz, 3H); ^13^C NMR (CDCl_­_, 125 MHz) *δ* 174.4, 70.3, 65.2, 63.3, 34.1, 31.9, 29.6-29.7 (7C), 29.58, 29.44, 29.35, 29.2, 29.1, 24.9, 22.7, 14.1; HR ESI-MS *m/z* 381.2984 [M + H]+ calcd for C_­_H_­_NaO_­_, 381.2975.

### Drug sensitivity test

Growth of yeast cells treated with purified aggreceride B, purchased fatty acids (Tokyo Chemical Industry), and synthesized monoacylglycerols was measured as described previously ^13^. Briefly, yeast cultures at mid-log phase were diluted to 0.0033 OD_­_and supplemented with varying concentrations of compounds. The culture conditions (incubation temperature, time, and medium) are described in figure legends, otherwise, cells were incubated in YES medium at 27°C for 24 hours. After incubation, the turbidity (OD_­_) was measured using a MULTISKAN FC (Thermo Scientific) or an Infinite F Plex (Tecan).

### Fluorescence microscopy

To observe the effect of fatty acids, 2 mg/mL DMSO solution of fatty acids were added to the mid-log phase culture with a final concentration of 10 µg/mL. Cells were incubated for three hours. Other conditions, if employed, are noted in the figure legends. For visualizing septa, a 1 mg/mL aqueous solution of calcofluor white (Fluorescent Brightener 28, Sigma-Aldrich) was added to the cell culture to a final concentration of 1 µg/mL. For detecting sterols in the PM, a 2 mg/mL DMSO solution of filipin complex (Sigma-Aldrich) was added to the culture medium to a final concentration of 10 µg/mL.

For the time-lapse observation of the growth of the septum, cells were attached to the bottom of 96-well Half Area High Content Imaging Glass Bottom Microplate (Corning). The glass bottoms of wells were first coated with lectin by spreading 1 µL of 1 mg/mL lectin solution and allowed to dry for at least 15 minutes with the lid on. Excess lectin was washed off with the medium used to cultivate cells. The cell culture diluted to 2.0 OD_­_was added to the well and left for five minutes after which the culture medium was removed from the well and gently washed with the medium to remove cells not adhering to the bottom. Finally, the medium was added to start the incubation.

To synchronize the cell cycle of *cdc25-22* temperature-sensitive mutant cells, cell cultures grown at 27°C to mid-log phase were transferred to 37°C and incubated for four hours to arrest the cell cycle. The cell cycle was resumed by rapidly shifting to 27°C and incubating at this temperature. Cells were chemically fixed and observed under fluorescence microscopy. 400 µL of the culture was mixed with 50 µL of 36% formalin and left for 30 minutes at room temperature. Cells were collected by centrifugation (2,000 rpm, 1 min.),washed three times by PBS, and then mounted.

BZ-X viewer software was used for image acquisition together with KEYENCE BZ-X710 fluorescence microscope equipped with a PlanApo 100x lens. Image processing and analyses were performed using ImageJ. Images whose background signals were removed are indicated in the figure legends. The intracellular distribution of Mid1-GFP and PM sterol were measured by line scan of the fluorescence intensity. The relative intensity of the fluorescence was calculated by taking the maximum intensity on each line profile as 100%. A z-stack consisting of nine images with 0.5 µm spacings was projected to a single image (maximum intensity projection), and the obtained image was used for analyzing the cortical distribution of Mid1-GFP.

### TEM observation

Yeast cells in mid-log phase were treated with DMSO or C15:0 for three hours in YES medium and harvested. Cells were fixed in a 2.5% glutaraldehyde 1×PBS (pH 7.5) solution for one hour. After washing with 1×PBS, they were further fixed in a 1% KMnO_­_solution and washed with distilled water. For dehydration, samples were suspended in 50% ethanol and left for ten minutes, then centrifuged at 3,000 rpm for five minutes to remove the supernatant, and the same procedure was repeated with 70%, 90%, 95%, and 99.5% ethanol. Next, the samples were washed with propylene oxide and left for five minutes. After centrifugation at 3,000 rpm for five minutes to remove the supernatant, 50% Spurr resin (Spurr resin: propylene oxide = 1:1) was added and left overnight. Then, the 50% Spurr resin was replaced with fresh 50% Spurr resin and left overnight. The Spurr resin was subsequently replaced with 100% Spurr resin, and propylene oxide was removed completely by leaving at reduced pressure in a desiccator for three hours. The samples were then left in an oven at 70°C for over ten hours. 100 nm-thin cross sections were obtained from the samples and were stained with 4% uranyl acetate for ten minutes and Reynold’s solution (0.4% lead citrate solution) for ten minutes. Electron microscopy was performed on a JEM-1010 transmission electron microscope (JEOL) with a 60 kV beam.

### Untargeted lipidomics

Wild-type and *erg31* Δ *erg32*Δ cells in mid-log phase were treated with DMSO or 10 µg/mL C15:0 for three or six hours at 27°C. Cells were harvested by centrifugation (2,000 rpm, 1 min.), and washed with ice-cold distilled water three times (2,000 rpm, 1 min.). 5×10^7^ cells were collected in tubes and flash-frozen in liquid nitrogen. Samples were stored at - 80°C before lipid extraction.

For the extraction of lipids, samples were first suspended in 100 µL methanol and sonicated for two minutes. To the cell suspension, 100 µL methanol containing internal standards (5 µL EquiSPLASH (Avanti Polar Lipids), 0.32 µL FA 16:0-d3 (10 mM), and 0.32 µL FA 18:0-d3 (10 mM)), 100 µL chloroform, and 10 µL distilled water were added. After further sonication for 2 min, samples were left at room temperature for two hours. Samples were centrifuged (5,000 rpm, 15 min) to collect supernatant. Lipids were subjected to an LC/Q-TOF-MS system and the obtained mass spectrometry data were analyzed with MS-DIAL4 ^28^ to conduct compound annotation. The experimental detail was described in our previous study ^49^

Hierarchical clustering and heatmap analysis were performed using the R package ‘pheatmap’. Log_­_-transformed quantitative data from untargeted lipidomics were used for the analyses. Euclidean distance was used to calculate the distances, and clustering was performed by Ward’s method.

### Real-time quantitative PCR

Wild-type and *Pnmt41-ole1* cells were harvested from a mid-log culture grown in EMM. Cells were washed with ice-cold distilled water and suspended in 0.4 mL of ISOGEN (Nippon Gene). 0.3 g of glass beads were added to the suspension and cells were disrupted with a Multi-Beads shocker (Yasui Kikai, 15 rounds of 30 sec. shake at 2500 rpm and 30 sec. interval). Total RNA was extracted according to the manufacturer’s protocol. The RNA concentration was measured by a DU730 UV/vis spectrophotometer (Beckman Coulter). cDNA was prepared with a PrimeScript RT reagent kit (TaKaRa). qPCR was performed on Analytikjena qTOWER3G. THUNDERBIRD Next SYBR (TOYOBO) was used for RT-PCR. Primer sequences are listed in **Extended Data File.** The expression of *ole1* was normalized to that of *act1*.

## Acknowledgements

We thank Drs. R. Fukuda and R. Iwama (Univ. Tokyo) for budding yeast strains, Drs. C. Boone (Univ of Toronto and RIKEN) and K. Kume (Hiroshima Univ.) for helpful discussions, Ms. A. Hori (RIKEN) for measuring yeast lipidome, and Dr. W. Fenical (SIO, UCSD) for supporting compound screen. A part of yeast strains was provided through the National BioResource Project – Yeast, Japan. The John Mung Program (Kyoto University) is acknowledged for the financial support to S.N. This work was supported in part by a Grant-in-Aid for Scientific Research from the Japan Society for the Promotion of Science (JP21H02128 to S.N.; JP19H05640 and JP23H05473 to S.N. and M.Y.; JP15H05897, JP15H05898, and JP20H00495 to M.A.; JP21K18216 and JP24K02011 to H.T.; JP17H06401, JP23H04882, and JP24H00493 to H.K), Transformative Research Areas (A) “Latent Chemical Space” from the Ministry of Education, Culture, Sports, Science and Technology, Japan (JP23H04880 and JP23H04882 to M.Y. and Y.Y.), JST National Bioscience Database Center (JPMJND2305, H.T.), RIKEN Pioneering Project “Glyco-Lipidologue Initiative” (M.A.), AMED Moonshot Research and Development Program grant number JP22zf0127007 (M.A.) and JST ERATO “Arita Lipidome Atlas Project” (JPMJER2101 to H.T. and M.A.). Y.H. was supported by the Japan Society for the Promotion of Science Research Fellowship for Young Scientists (JP21J21126).

## Author contributions

Y.H., M.Y., and S.N. designed the research and analyzed the data; Y.H. conducted most experiments; N.S. conducted experiments using mutants overexpressing Lcf1 or Lcf2; A.M. and Y.Y. prepared genetic materials; S.K. supported TEM analysis; H.T. and M.A conducted lipidomics; S.N. conducted compound screen with the help of H.K. and R.I.; Y.H. and S.N. wrote the draft manuscript; Y.H., H.T., A.M., Y.Y., M.A., M.Y., and S.N. edited the manuscript.

## Competing interests

The authors declare no competing interests.

---

**Extended Data Figure S1.**
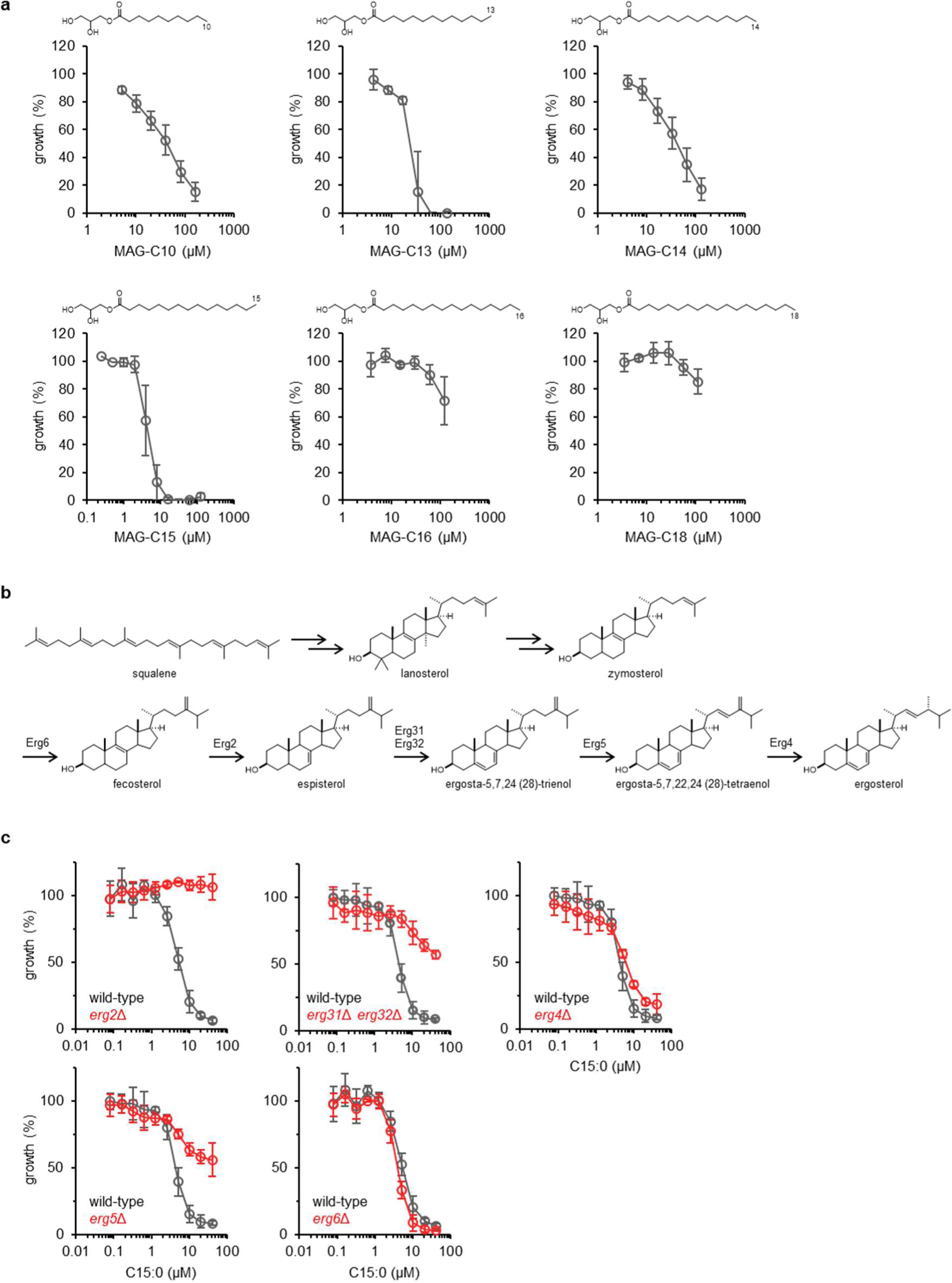
The activity of monoacylglycerols and C15:0. **a,** Monoacylglycerol sensitivity of wild-type cells. Monoacylglycerols with C10:0 (MAG-C10), C13:0 (MAG-C13), C14:0 (MAG-C14), C15:0 (MAG-C15), C16:0 (MAG-C16) and C18:0 (MAG-C18) were tested. MAG-C10, MAG-C13, MAG=C16, MAG-C18, n = 4; MAG-C14, n = 5; MAG-C15, n = 3. Error bar = SD. **b,** Ergosterol biosynthetic pathway. **c,** C15:0 sensitivity of *erg2*Δ, *erg4*Δ, *erg5*Δ, *erg6*Δ mutants. Grey and red lines represent wild-type cells and *erg* mutants, respectively. n = 3. Error bar = SD.

**Extended Data Figure S2.**
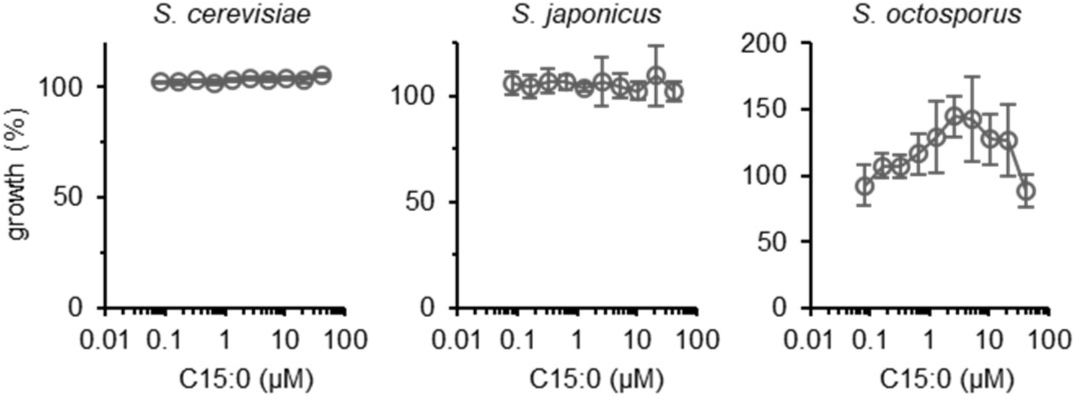
C15:0 does not affect the growth of yeast species other than *S. pombe*. C15:0 sensitivity of the budding yeast *S. cerevisiae* and the fission yeasts *S. japonicus* and *S. octosporus. S. cerevisiae* was cultured in the YPD medium, and *S. japonicus* and *S. octosporus* in the YES medium. The cultures were incubated at 27°C for 24 h. n = 3. Error Bar = SD.

**Extended Data Figure S3.**
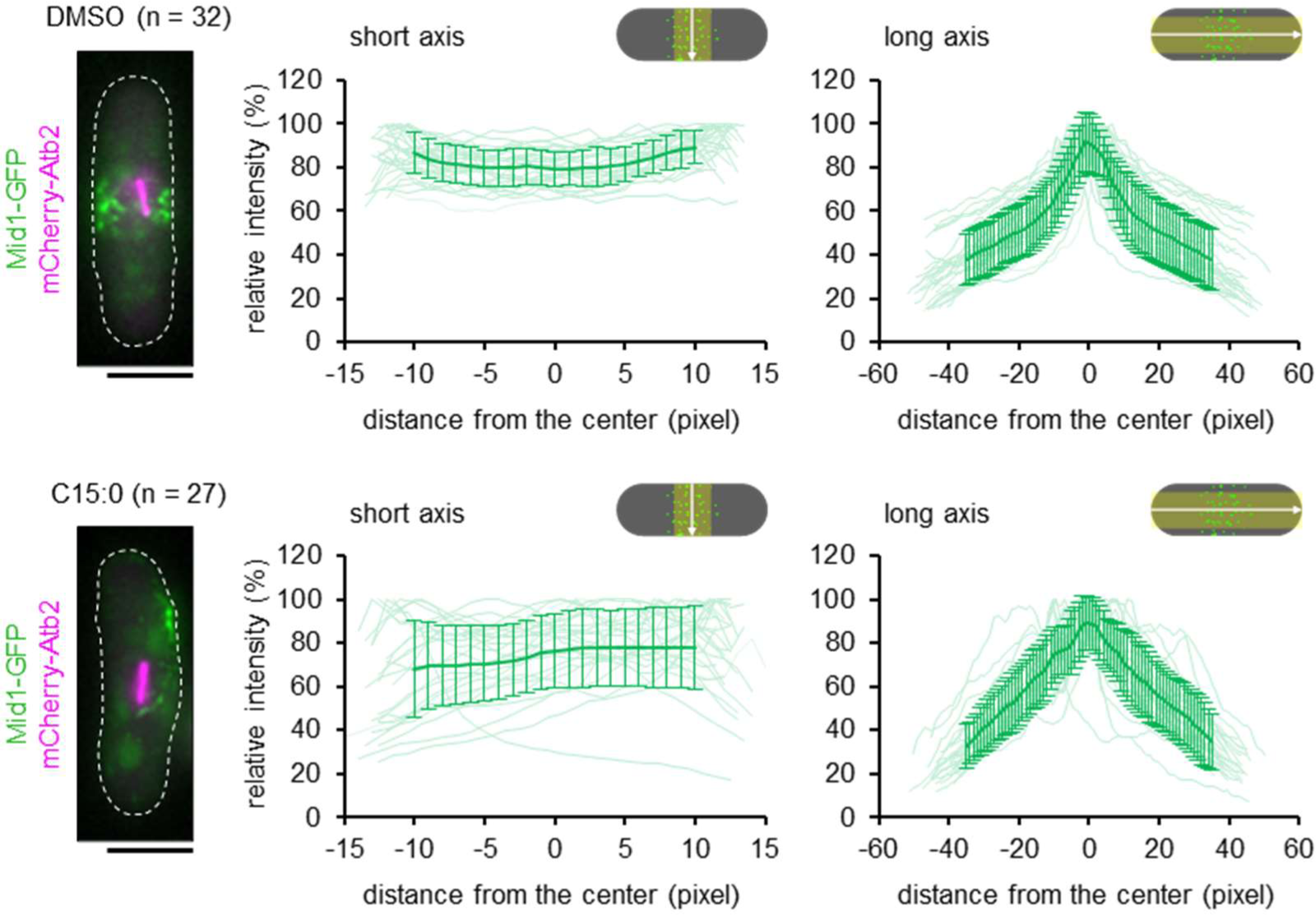
C15:0 disturbs the distribution of Mid1 in early mitotic cells. Distributions of Mid1-GFP in early mitotic cells. The stage of mitosis was determined by microtubule marked by mCherry-Atb2. The image on the left shows the representative cell for each condition. Mid1-GFP intensity was measured across a line with a 30-pixel width along the short and long axis of the cell. Each faint line in the graph is the result from a different cell, and the bold line shows the average. Error bar = SD. Scale bar = 5 µm.

**Extended Data Figure S4.**
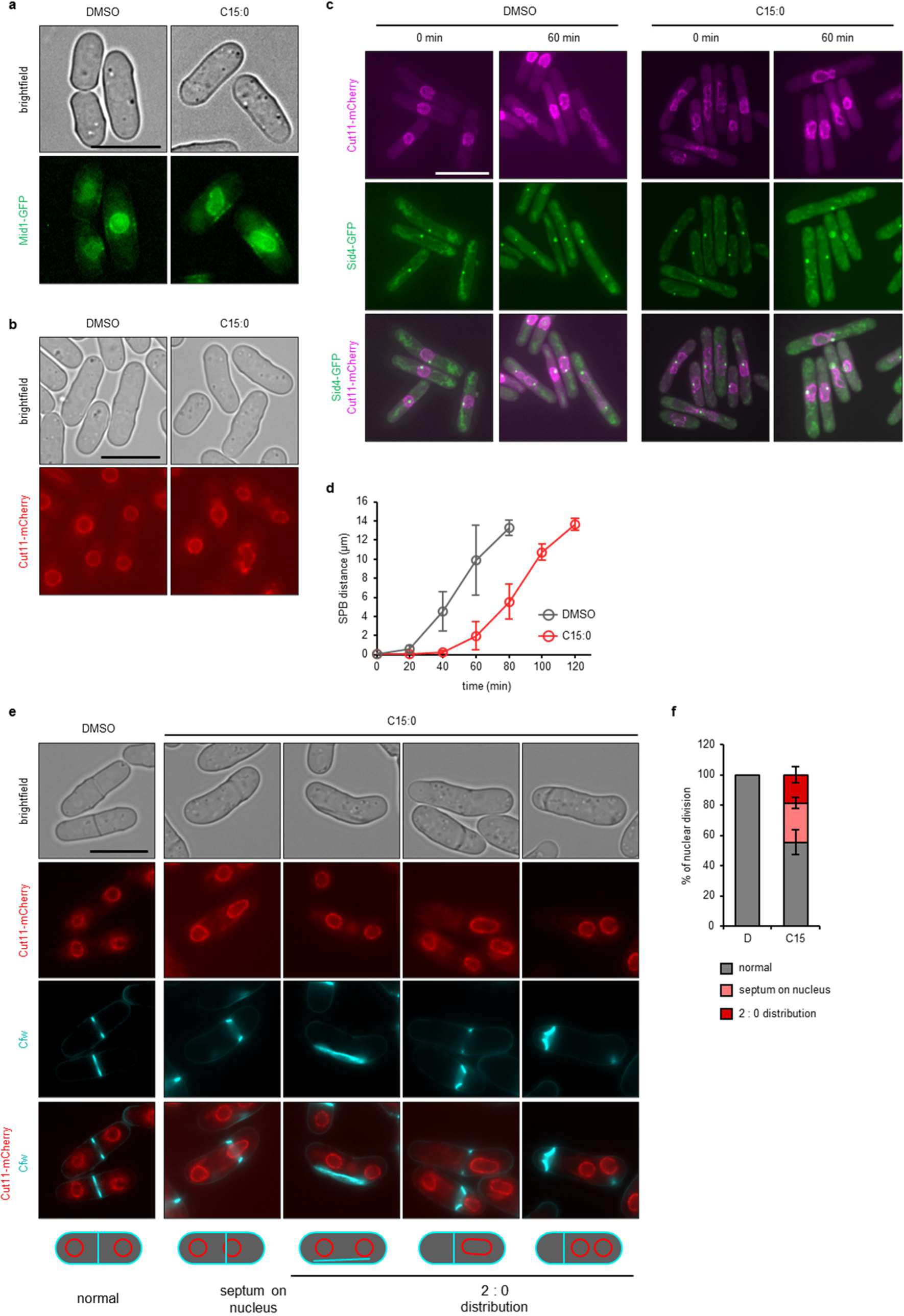
C15:0 affects nuclear morphology and mitosis. **a,** Nuclei marked by Mid1-GFP in interphase cells. The background signal was removed using ImageJ. Scale bar = 10 µm。 **b,** Nuclear envelope marked by Cut11-mCherry. Scale bar = 10 µm. **c,** SPB (Sid4-GFP) and nuclear envelope (Cut11-mCherry) in *cdc25-22* cells. Cells incubated at a restrictive temperature (27°C) were transitioned to a permissive temperature (27°C) to synchronize the cell cycle and induce mitosis. DMSO or C15:0 was added to the culture when incubation at 37°C was initiated. The observation was performed at indicated time points. SPB segregation was observed at 60 min in the control condition, but not in the majority of the C15:0 treated cells. **d,** Timelapse measurement of SPB distance. *cdc25-22* cells were cultured under the same conditions as in (**c**). The distance between segregated SPBs was measured after the transition to permissive temperature. n > 15 cells from three independent experiments. Error bar = SD. **e,** Observation of the nuclear envelope (Cut11-mCherry) and the septum (Cfw). Cartoons depict nuclear division patterns. Scale bar = 10 µm. **f,** Frequency of each nuclear division pattern. Cells were treated with DMSO or C15:0 for six hours. n > 40 cells with septa from three independent experiments. Error bar = SD.

**Extended Data Figure S5.**
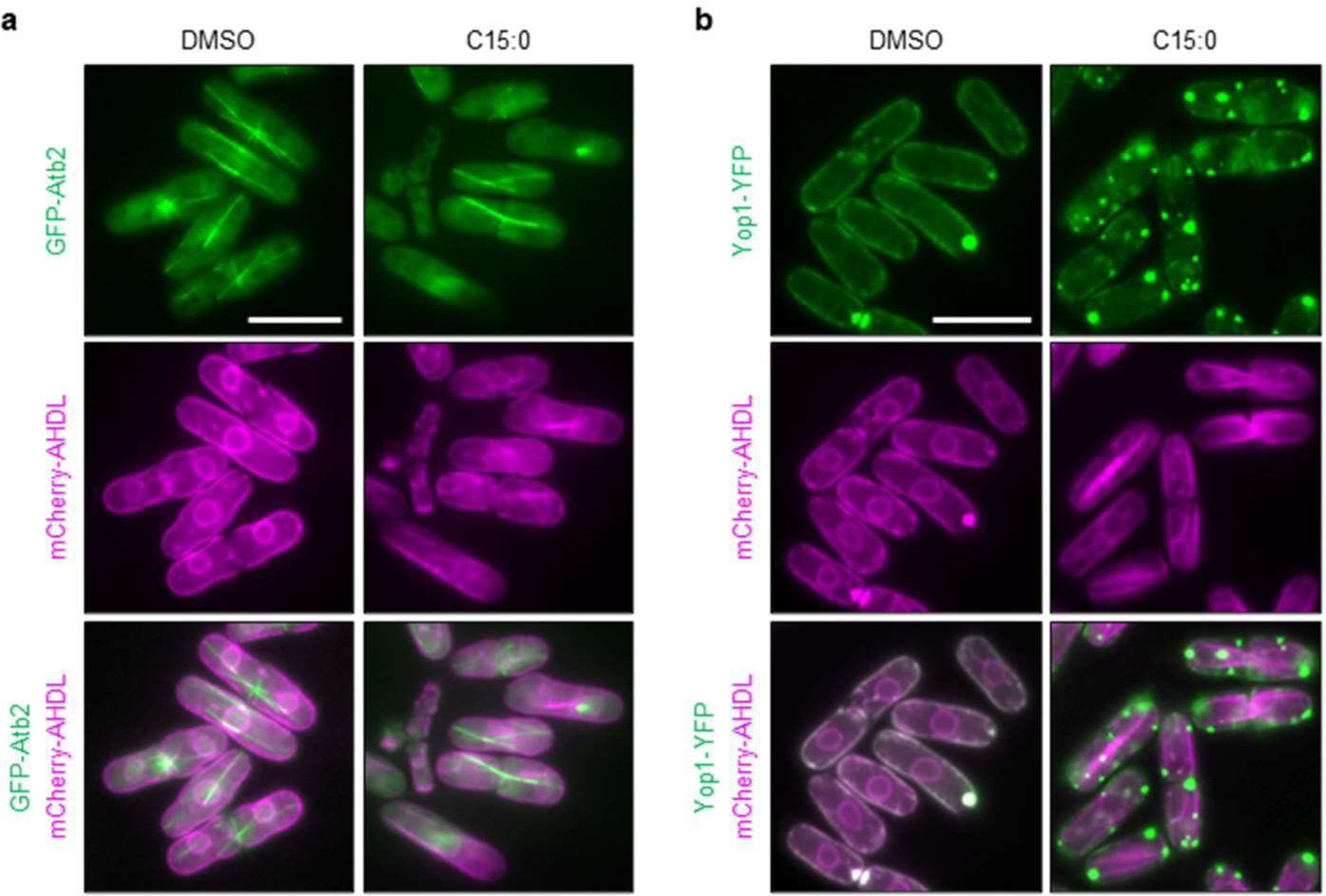
C15:0 induces the formation of planar ER architectures. **a,** Observation of the ER (mCherry-AHDL) and the microtubule. Scale bar = 10 µm。 **b,** Localization of YFP-tagged Yop1 (a tubule-forming protein). mCherry-AHDL was coexpressed to visualize the ER. Scale bar = 10 µm. Yop1-YFP was expressed from pDUAL constructs ^47, 50^. Scale bar = 10 µm.

**Extended Data Figure S6.**
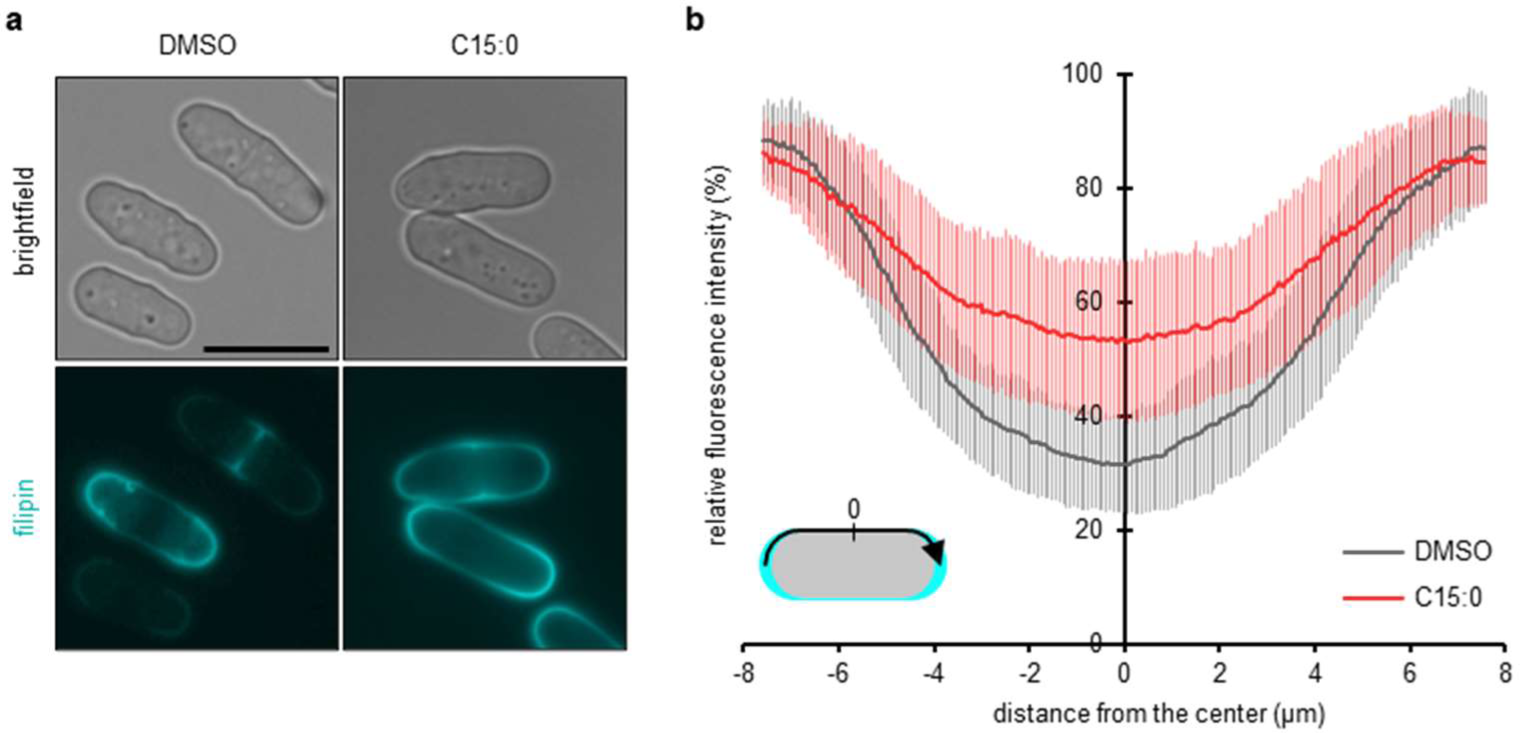
C15:0 attenuates the polarity of the PM sterols. **a,** Plasma membrane sterols visualized by filipin. Cells were treated with DMSO (control) or C15:0 for 6 hours. Scale bar = 10 µm. **b,** Distribution of plasma membrane sterols in interphase cells. The filipin intensity was measured along the cell periphery from one tip to the other. n > 90 cells from three independent experiments. Error bar = SD.

**Extended Data Figure S7.**
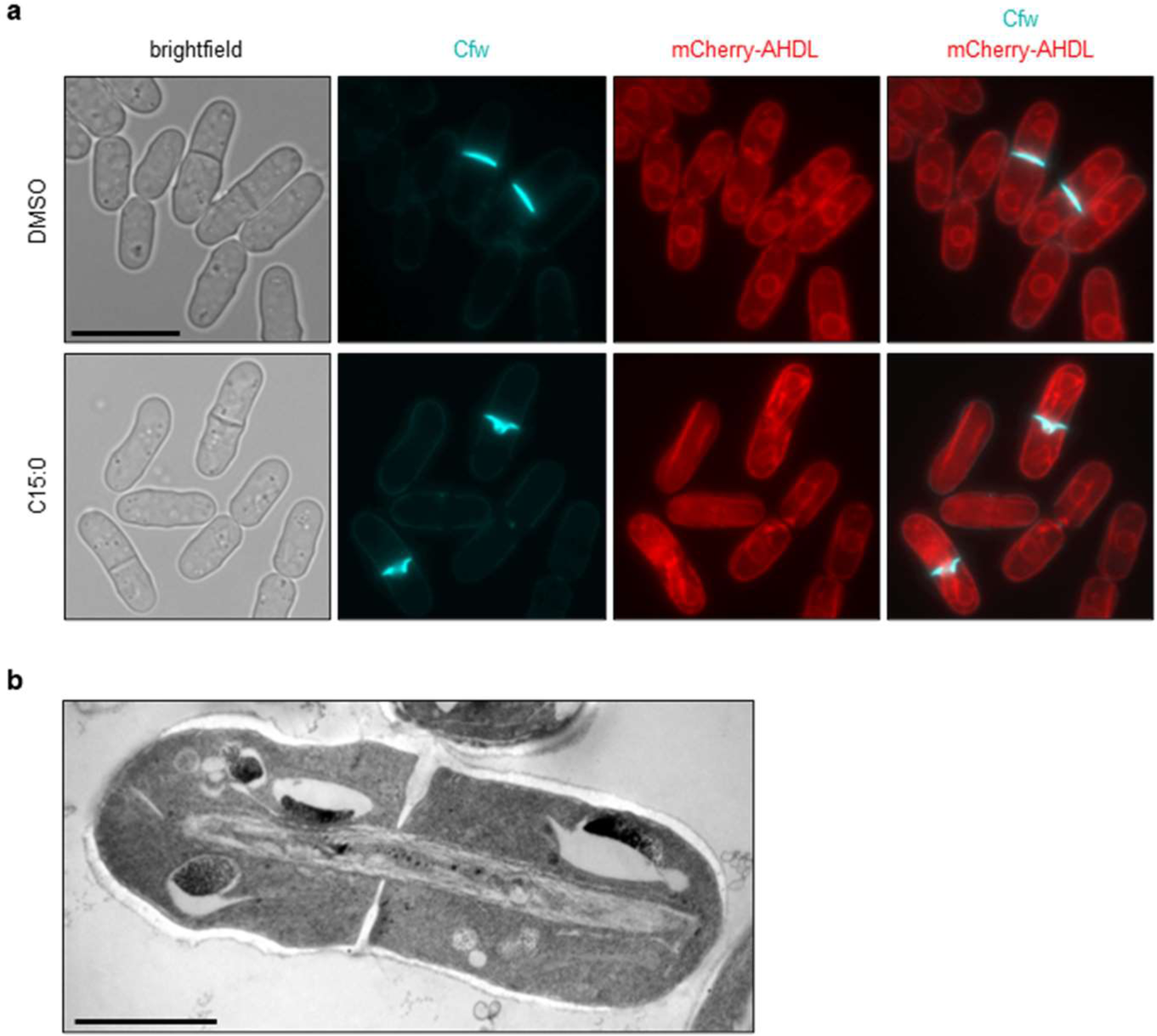
The giant ER sheet physically blocks the septal growth. **a,** Simultaneous observation of the septum and the ER. Scale bar = 10 µm. **b,** The rod-like structure of the ER observed in some C15:0 treated cells. Scale bar = 2 µm.

**Extended Data Figure S8.**
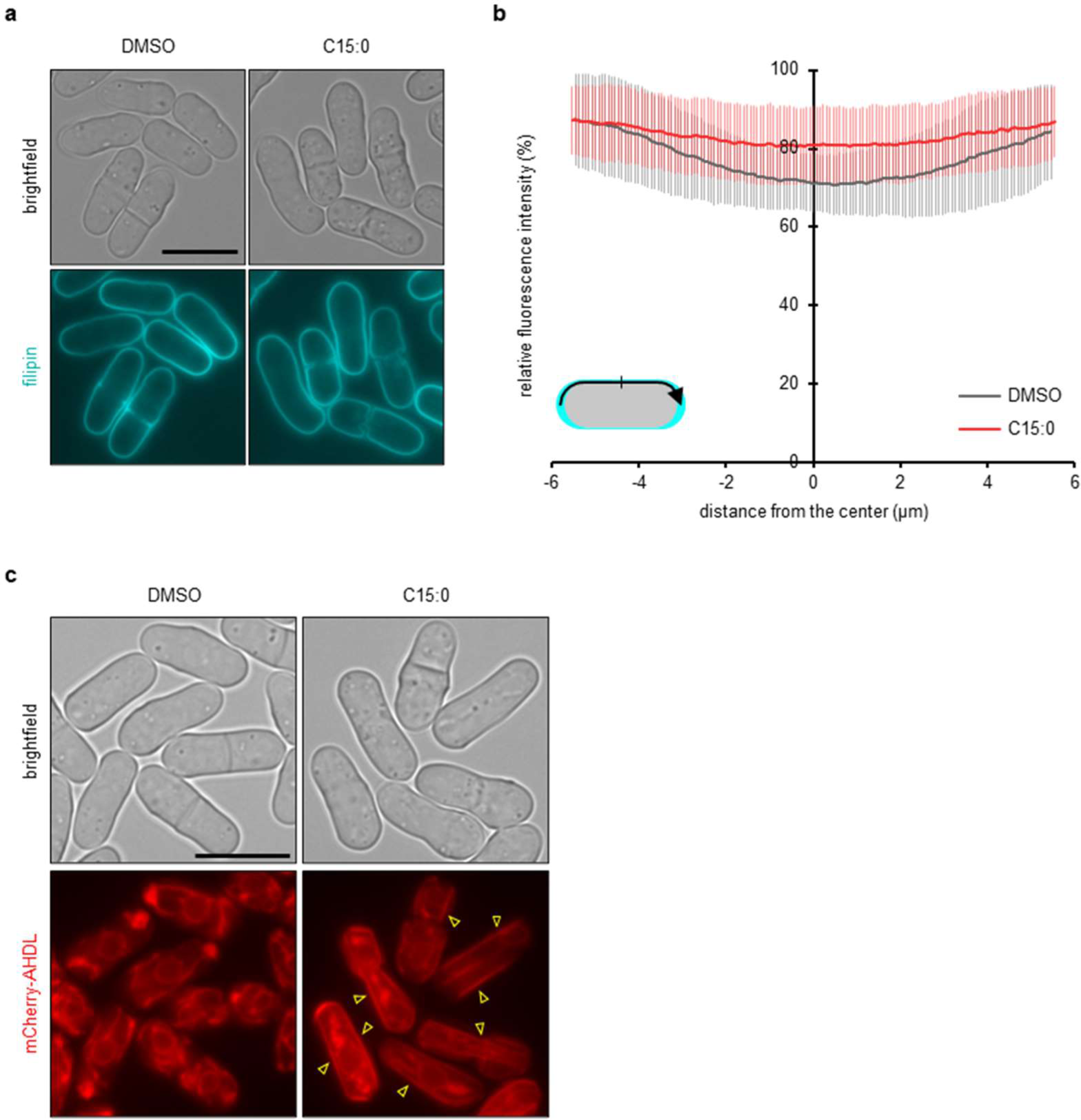
*scs*2Δ *scs*22Δ cells deficient in ER-PM contact exhibit disparate C15:0-induced phenotypes. **a,** Plasma membrane sterols visualized by filipin in *scs2*Δ *scs22*Δ cells. Cells were treated with DMSO (control) or C15:0 for 3 hours. Scale bar = 10 µm. **b,** Distribution of plasma membrane sterols in interphase *scs2*Δ *scs22*Δ cells. The filipin intensity was measured along the cell periphery from one tip to the other. n > 90 cells from three independent experiments. Error bar = SD. **c,** The ER in *scs2*Δ *scs22*Δ cells. Scale bar = 10 µm.

**Extended Data Figure S9.**
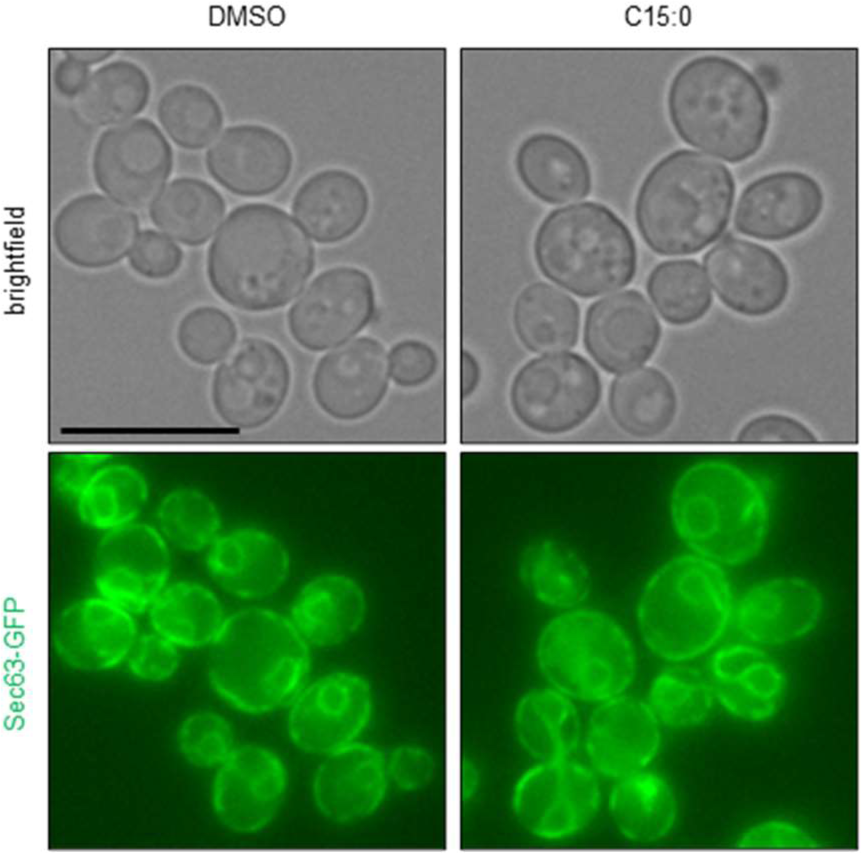
C15:0 does not affect the ER morphology in *S. cerevisiae* cells. Sec63-GFP was used for the marker to visualize the ER. Scale bar = 10 µm.

**Extended Data Figure 10.**
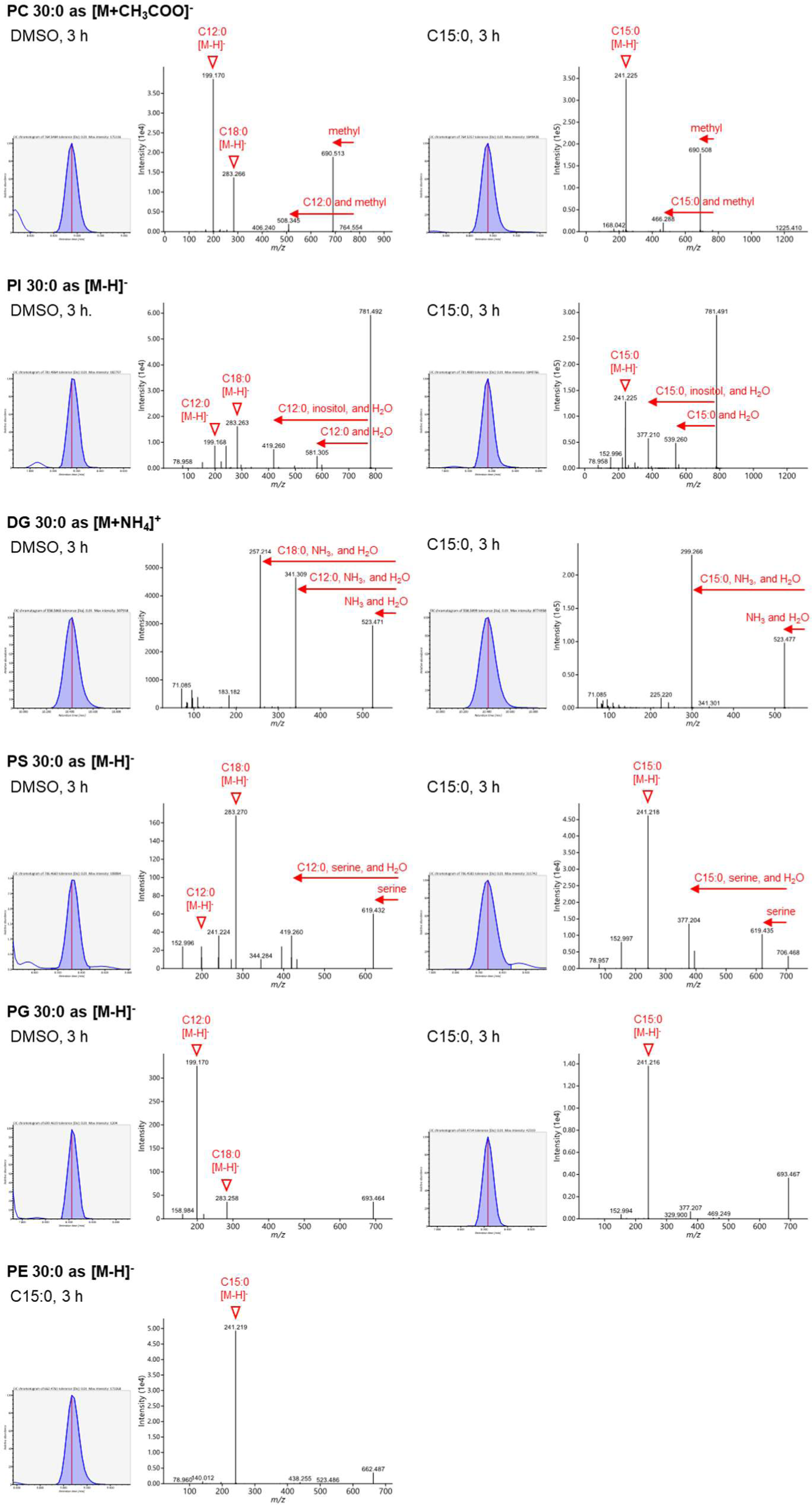
MS chromatograms and MS/MS spectra for GPLs 30:0. MS chromatograms and MS/MS spectra for GPLs (30:0) under DMSO or C15:0 treatment for three hours are shown. The intensity of the MS signals in the MS/MS spectrum for PE 30:0 in the DMSO-treated samples did not have enough intensity.

**Extended Data Figure S11.**
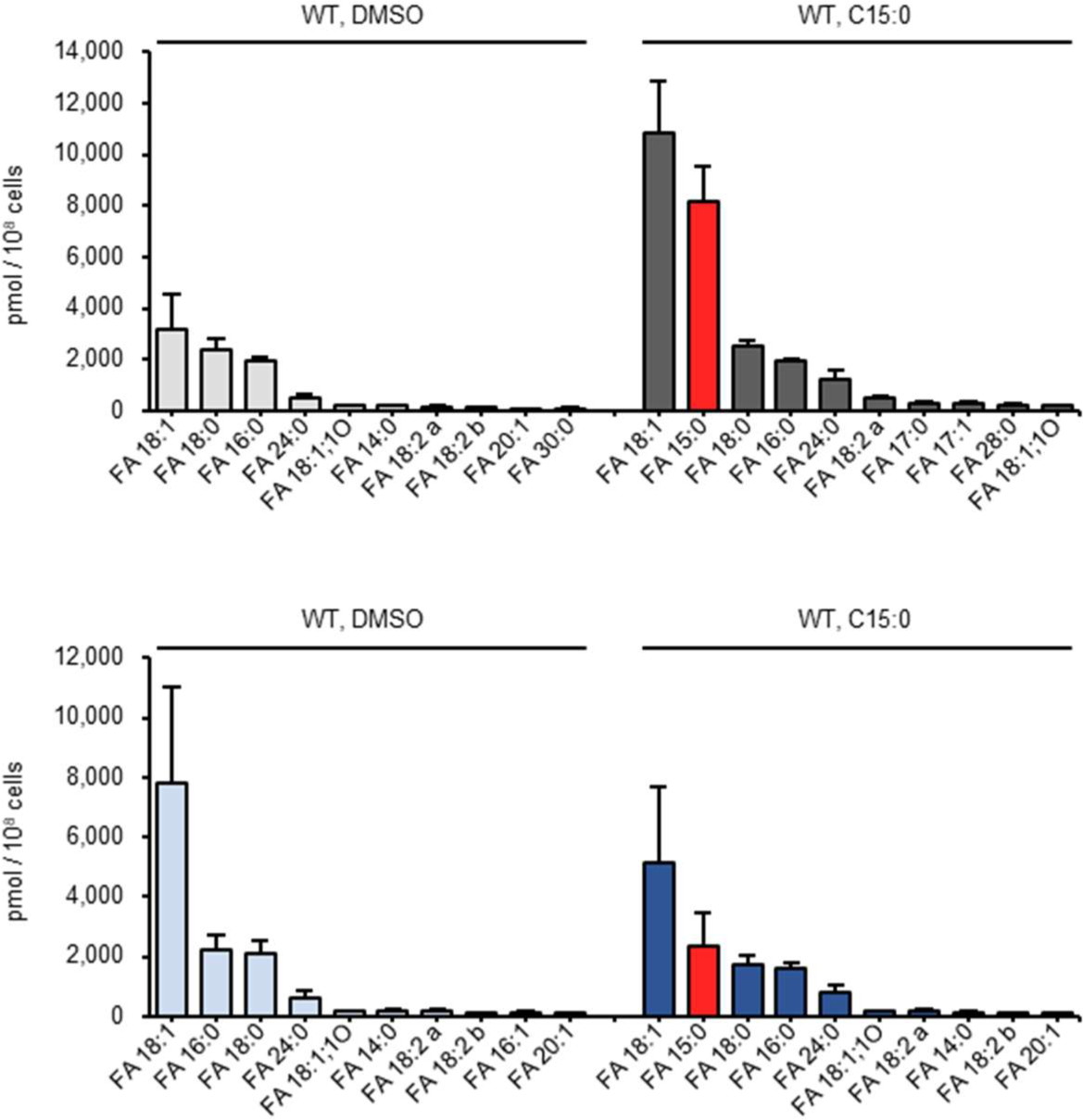
Profile of free fatty acids at six-hour chemical treatment. The quantity (pmol/10^8^ cells) of top ten molecular species under indicated conditions are shown. *erg* means *erg31* Δ *erg32* Δ mutant.

**Extended Data Figure S12.**
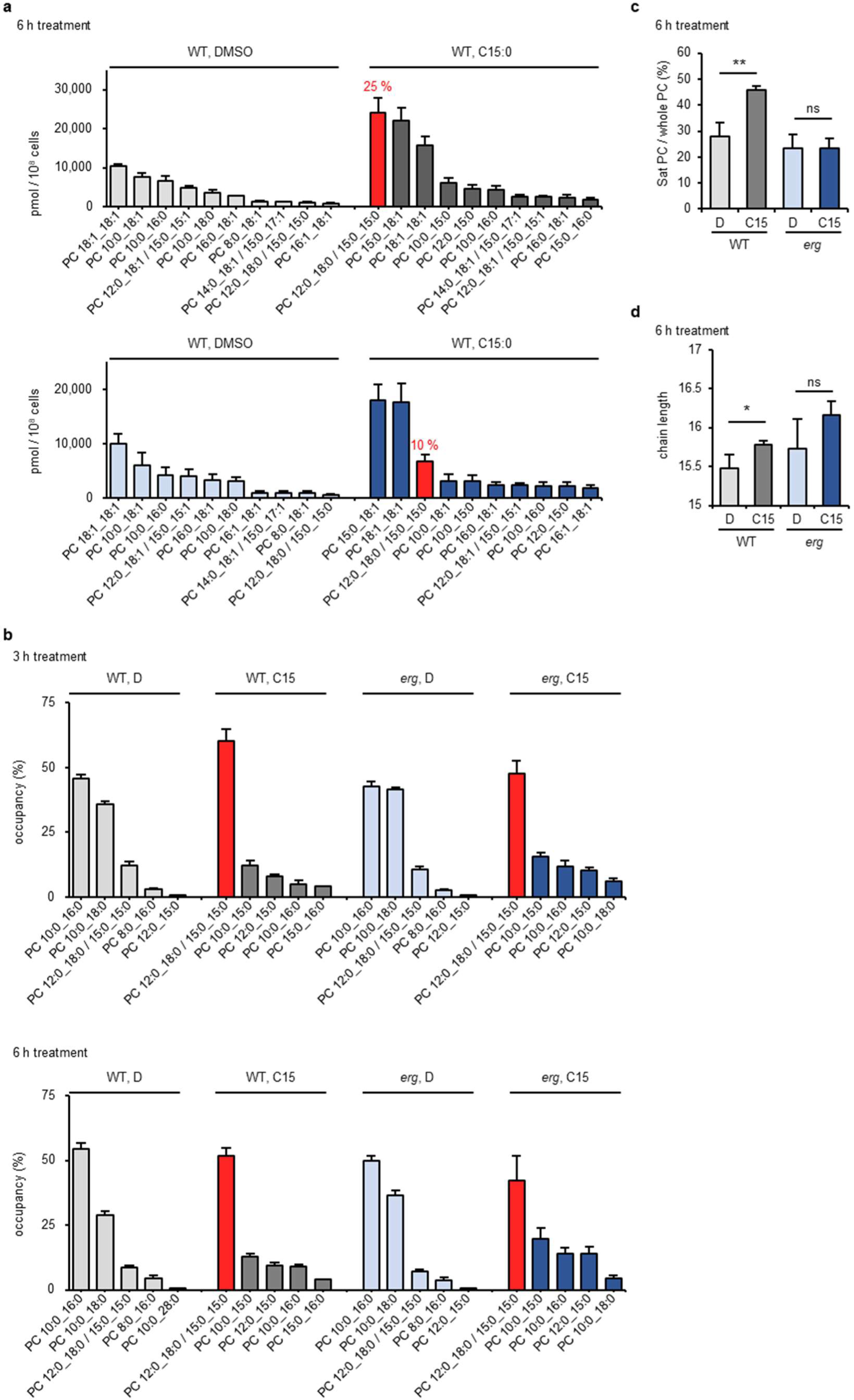
Effect of C15:0 on the profile of PC. **a,** Profile of PC at six-hour chemical treatment. The quantity (pmol/10^8^ cells) of top ten molecular species under indicated conditions are shown. The red bar indicates the species possessing two C15:0 as acyl chains and its proportion (%) is shown. PC 12:0_18:0 and PC 15:0_15:0 are shown together due to the limitations at LC separation. **b,** Composition of saturated PC. The proportion (%) of top five molecular species under indicated conditions are shown. The red bar indicates the species possessing two C15:0 as acyl chains. (**c**) The proportion (%) of saturated PC among total PC at six-hour chemical treatment. **d,** The average carbon chain length of acyl chains in PC at three-hour chemical treatment. (**a-d**) *erg* means *erg31*Δ *erg32*Δ mutant. (**a-d**) n = 4. Error bar = SD。 (**c-d**) P-values determined by unpaired two-tailed t-test. ns, no significance; * p < 0.05; ** p < 0.01; *** p < 0.001.

**Extended Data Figure S13.**
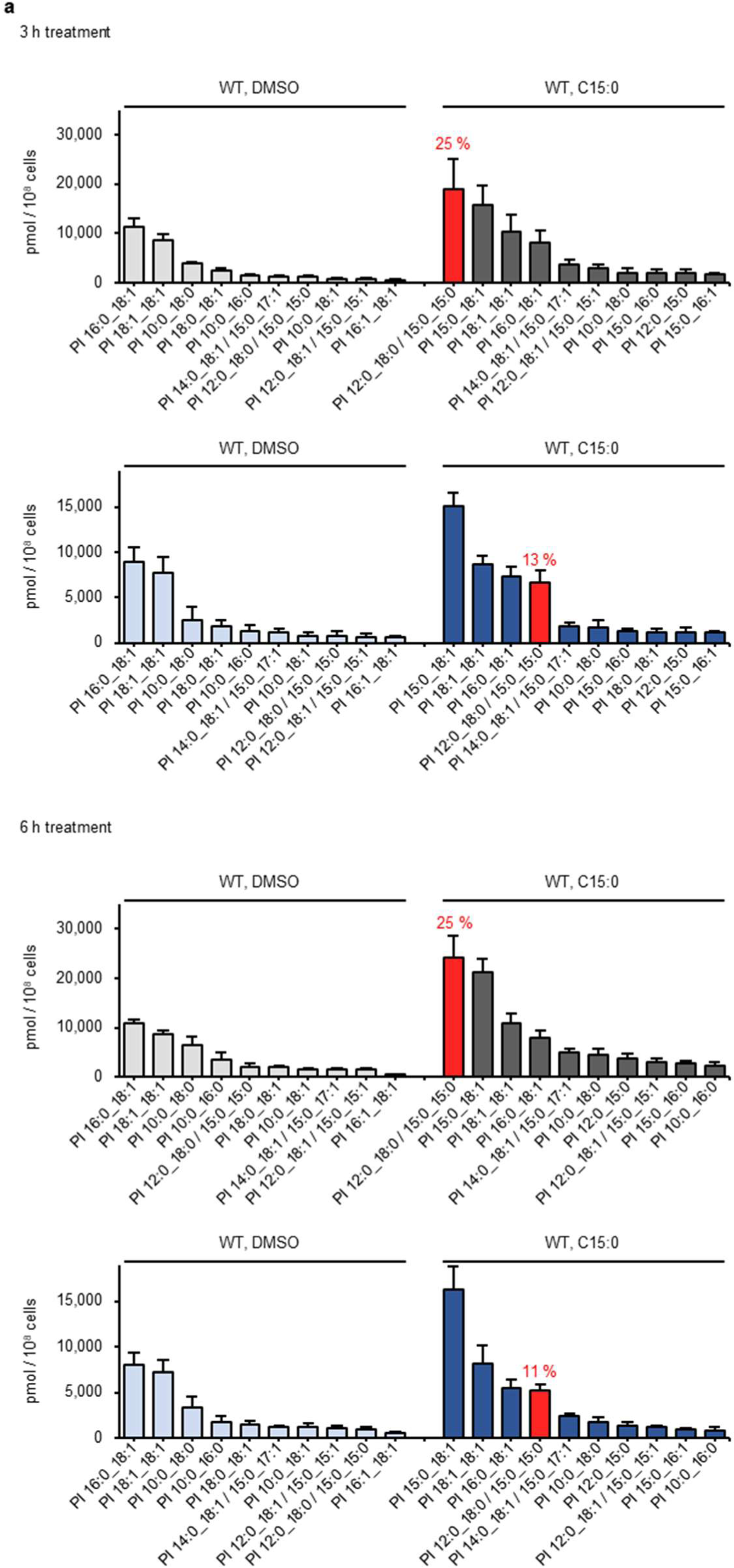
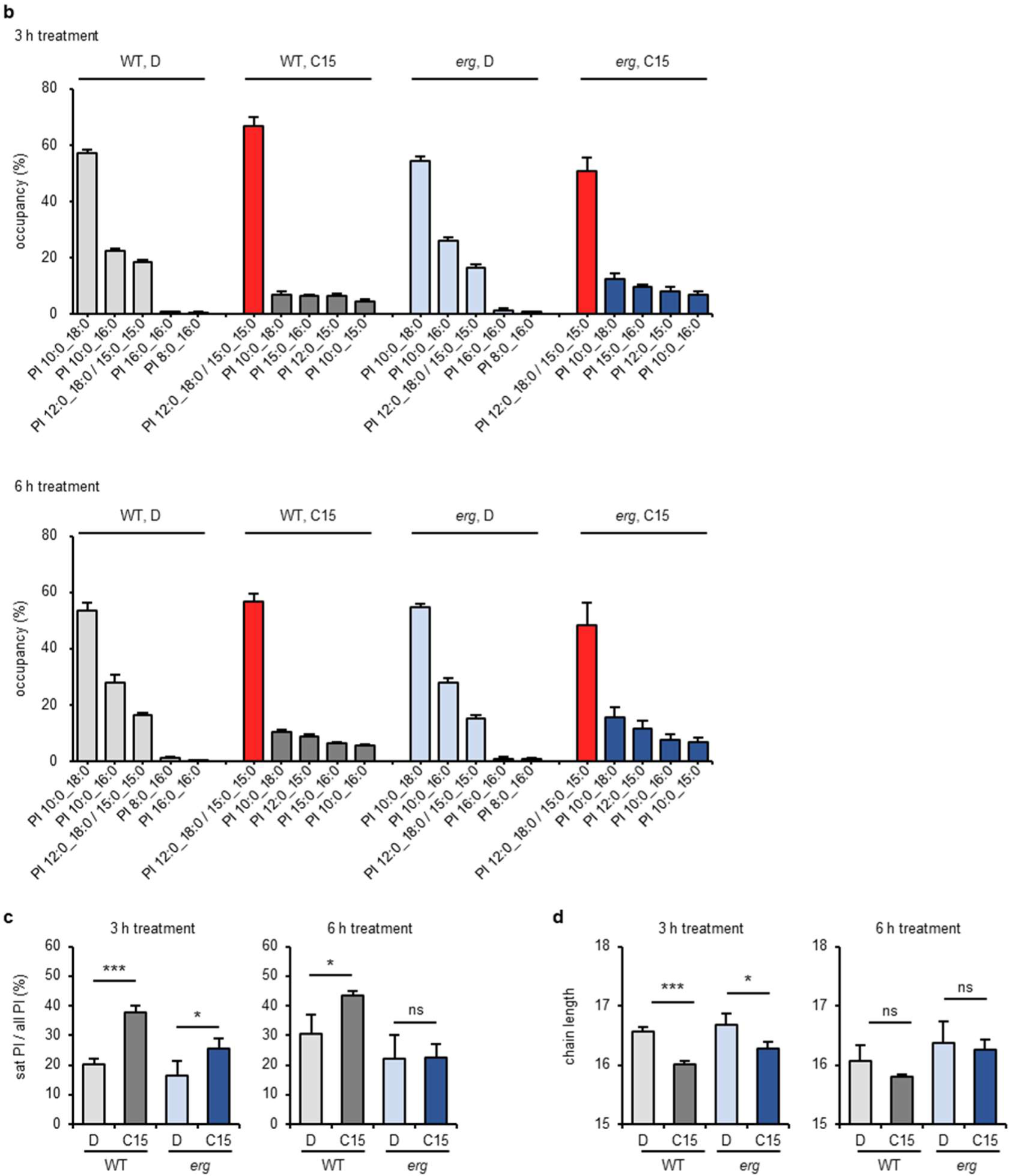
Effect of C15:0 on the profile of PI. **a,** Profile of PI at three or six-hour chemical treatment. The quantity (pmol/10^8^ cells) of top ten molecular species under indicated conditions are shown. The red bar indicates the species possessing two C15:0 as acyl chains and its proportion (%) is shown. The species that were not distinguished in LC are shown together. The species that were not distinguished at LC are shown together. **b,** Composition of saturated PI. The proportion (%) of top five molecular species under indicated conditions are shown. The red bar indicates the species possessing two C15:0 as acyl chains. **c,** The proportion (%) of saturated PI among total PI. **d,** The average carbon chain length of acyl chains in PI. (**a-d**) *erg* means *erg31*Δ *erg32*Δ mutant. (**a-d**) n = 4. Error bar = SD。 (**c-d**) P-values determined by unpaired two-tailed t-test. ns, no significance; * p < 0.05; ** p < 0.01; *** p < 0.001.

**Extended Data Figure S14.**
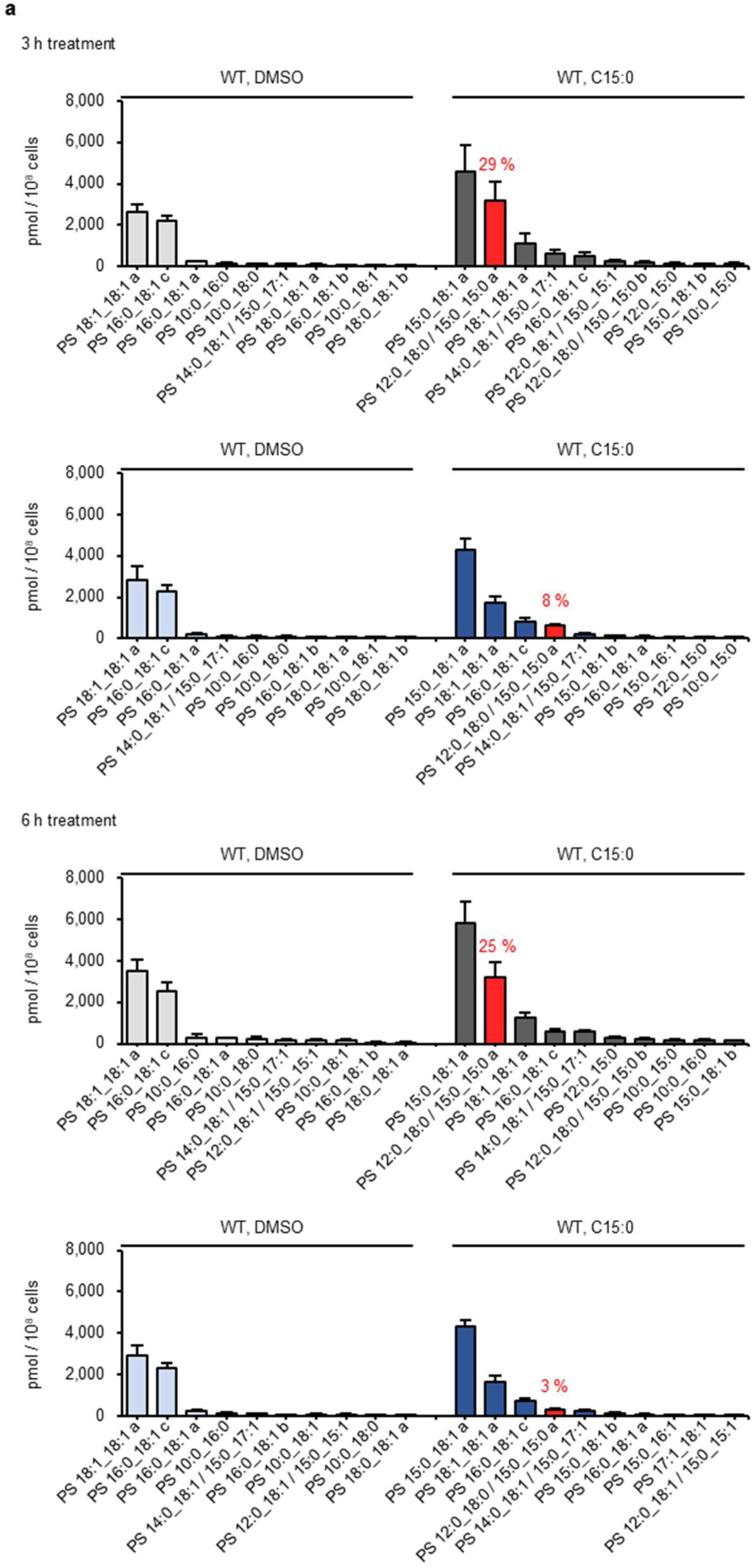
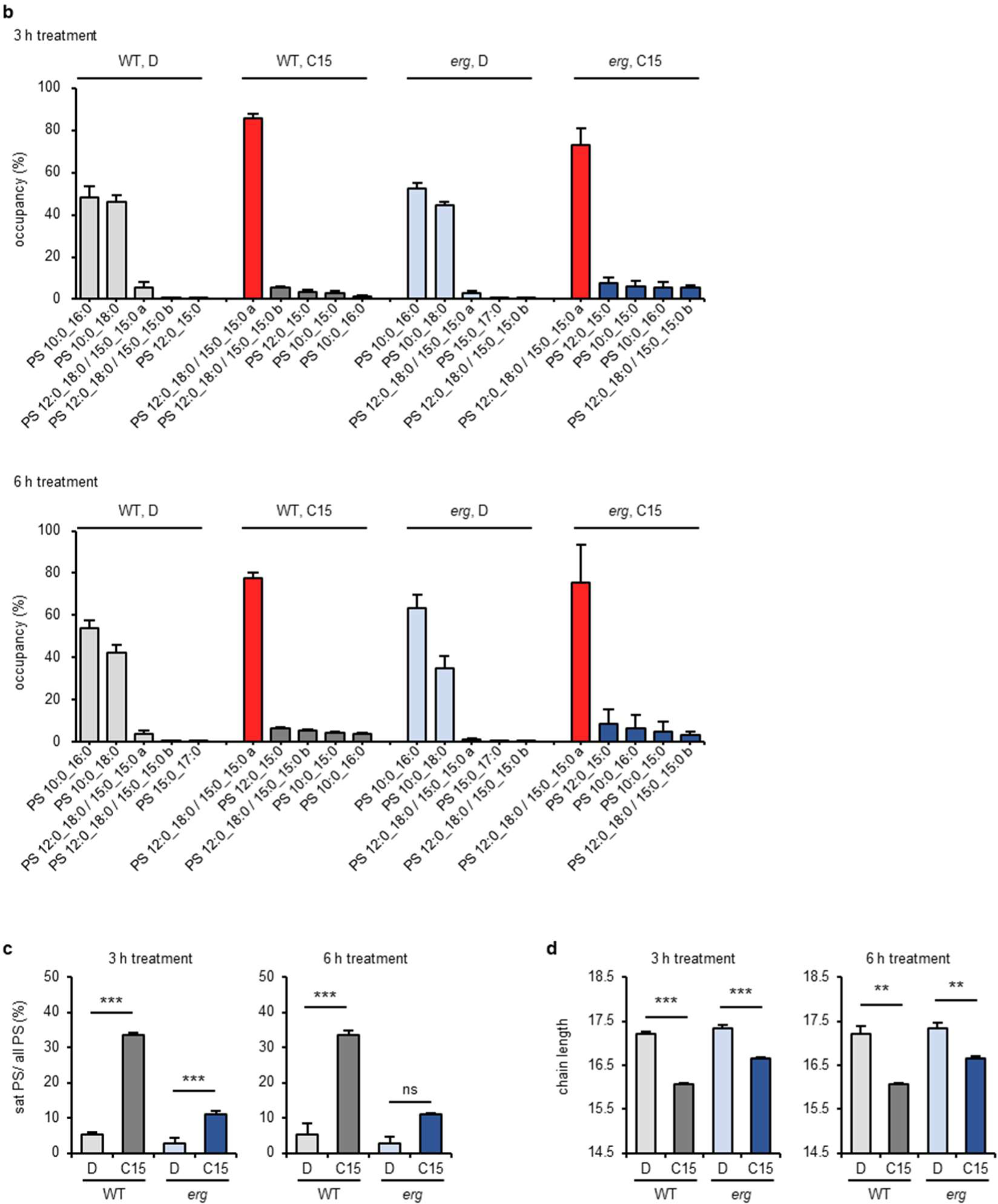
Effect of C15:0 on the profile of PS. **a,** Profile of PS at three or six-hour chemical treatment. The quantity (pmol/10^8^ cells) of top ten molecular species under indicated conditions are shown. The red bar indicates the species possessing two C15:0 as acyl chains and its proportion (%) is shown. The species that were not distinguished at LC are shown together. Note that molecules detected separately at LC but annotated to have the same structures are distinguished by alphabetical codes. **b,** Composition of saturated PS. The proportion (%) of top five molecular species under indicated conditions are shown. The red bar indicates the species possessing two C15:0 as acyl chains. **c,** The proportion (%) of saturated PS among total PS. **d,** The average carbon chain length of acyl chains in PS. (**a-d**) *erg* means *erg31*Δ *erg32*Δ mutant. (**a-d**) n = 4. Error bar = SD。 (**c-d**) P-values determined by unpaired two-tailed t-test. ns, no significance; * p < 0.05; ** p < 0.01; *** p < 0.001.

**Extended Data Figure S15.**
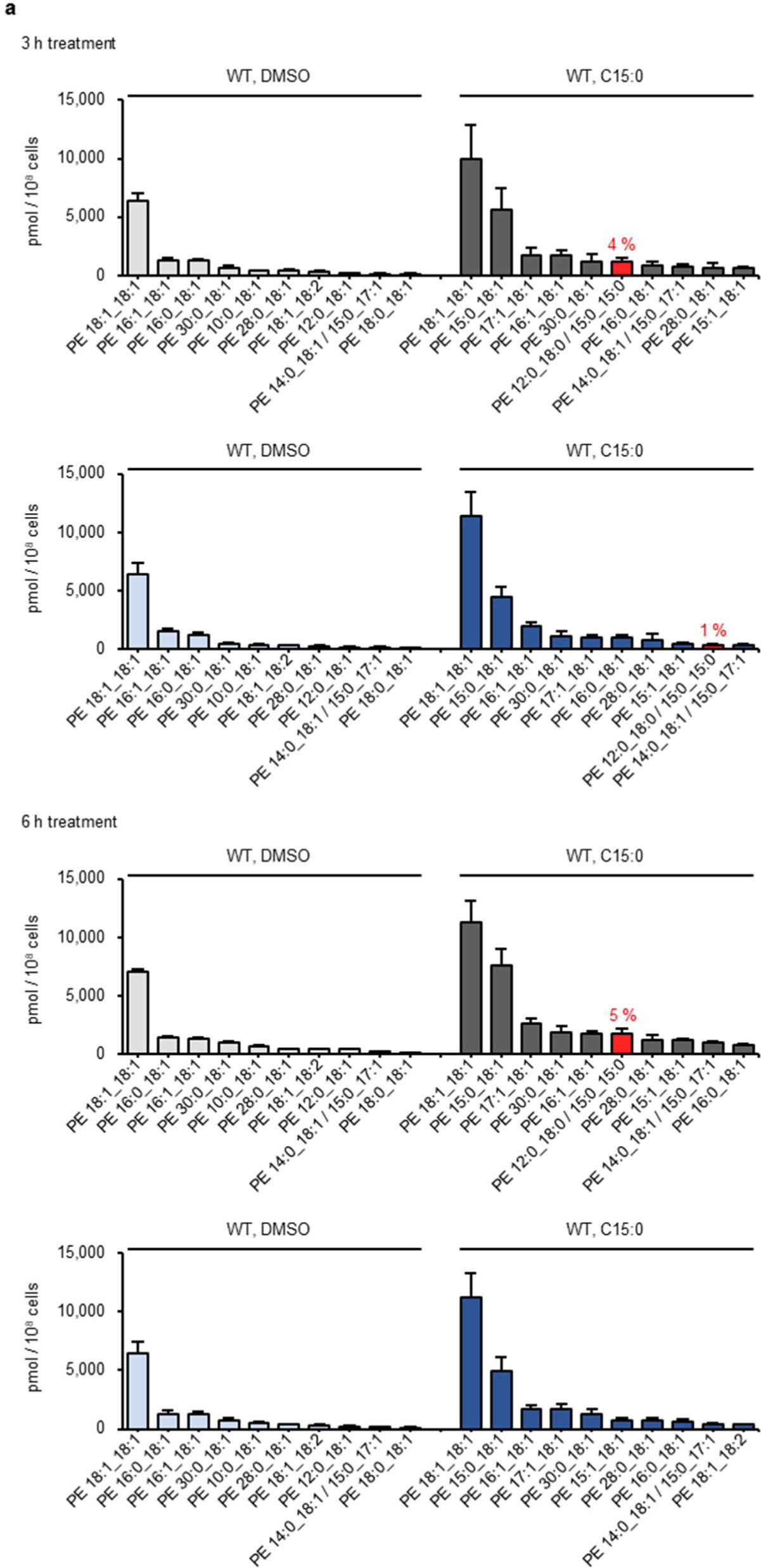
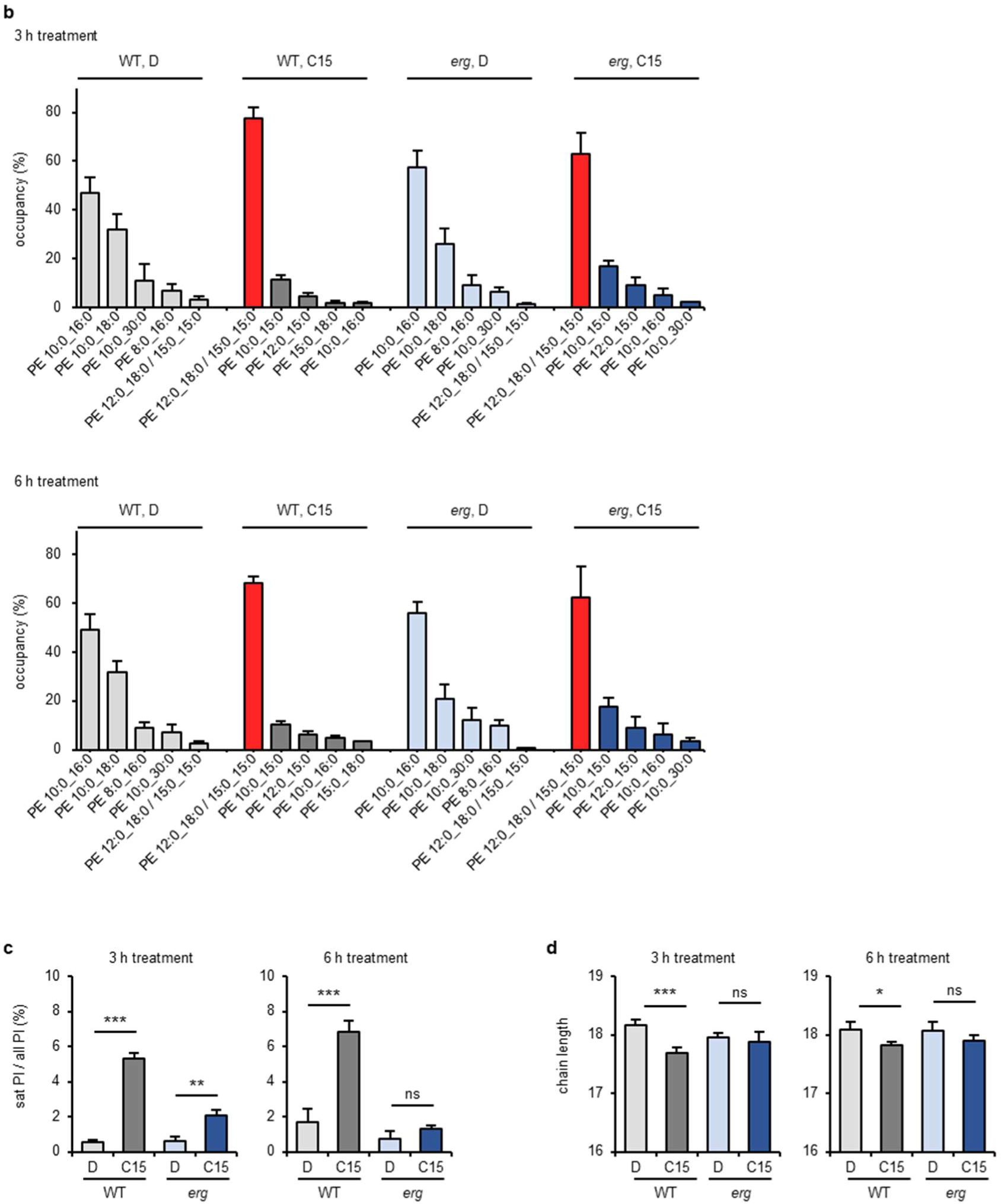
Effect of C15:0 on the profile of PE. **a,** Profile of PE at three or six-hour chemical treatment. The quantity (pmol/10^8^ cells) of top ten molecular species under indicated conditions are shown. The red bar indicates the species possessing two C15:0 as acyl chains and its proportion (%) is shown. The species that were not distinguished at LC are shown together. **b,** Composition of saturated PE. The proportion (%) of top five molecular species under indicated conditions are shown. The red bar indicates the species possessing two C15:0 as acyl chains. **c,** The proportion (%) of saturated PE among total PE. **d,** The average carbon chain length of acyl chains in PE. (**a-d**) *erg* means *erg31*Δ *erg32*Δ mutant. (**a-d**) n = 4. Error bar = SD。 (**c-d**) P-values determined by unpaired two-tailed t-test. ns, no significance; * p < 0.05; ** p < 0.01; *** p < 0.001.

**Extended Data Figure S16.**
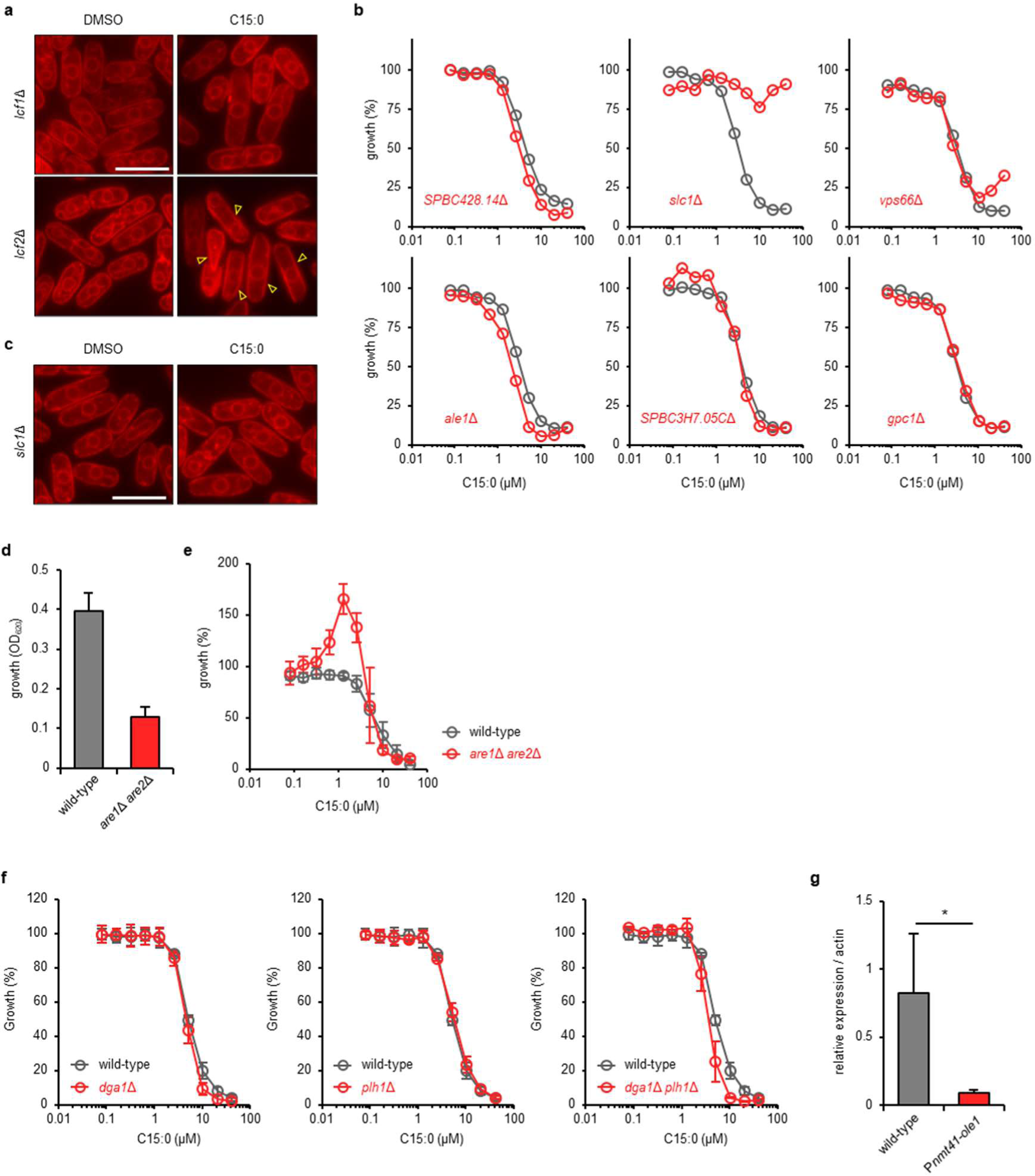
C15:0 is incorporated into membrane lipids to exert its activity. **a,** The ER in *lcf1*Δ and *lcf2*Δ cells. Scale bar = 10 µm. **b,** C15:0 sensitivity of *SPBC428.14* Δ, *slc1*Δ, *vps66* Δ, *ale1* Δ, *SPBC3H7.05C* Δ, and *gpc1* Δ cells. The strains from the Bioneer gene deletion library were used. Grey and red lines represent wild-type and mutant cells, respectively. n = 2. **c,** The ER in *slc1*Δ cells. Scale bar = 10 µm. **d,** Growth of *are1*Δ *are2*Δ cells. OD595 of the cultures after 24 hours of incubation under control conditions without chemical treatment in the C15:0 sensitivity test are shown in the graph. n = 3. Error bar = SD. **e,** C15:0 sensitivity of *are1*Δ *are2*Δ cells. n = 3. Error bar = SD. **f,** C15:0 sensitivity of *dga1* Δ, *plh1* Δ, and *dga1* Δ *plh1* Δ cells. n = 3; Error bar = SD. **g,** *ole1* mRNA level in wild-type and P*nmt41-ole1* cells. n = 4; Error bar = SD. P-values determined by unpaired two-tailed t-test. ns, no significance; * p < 0.05.

**Extended Data Figure S17.**
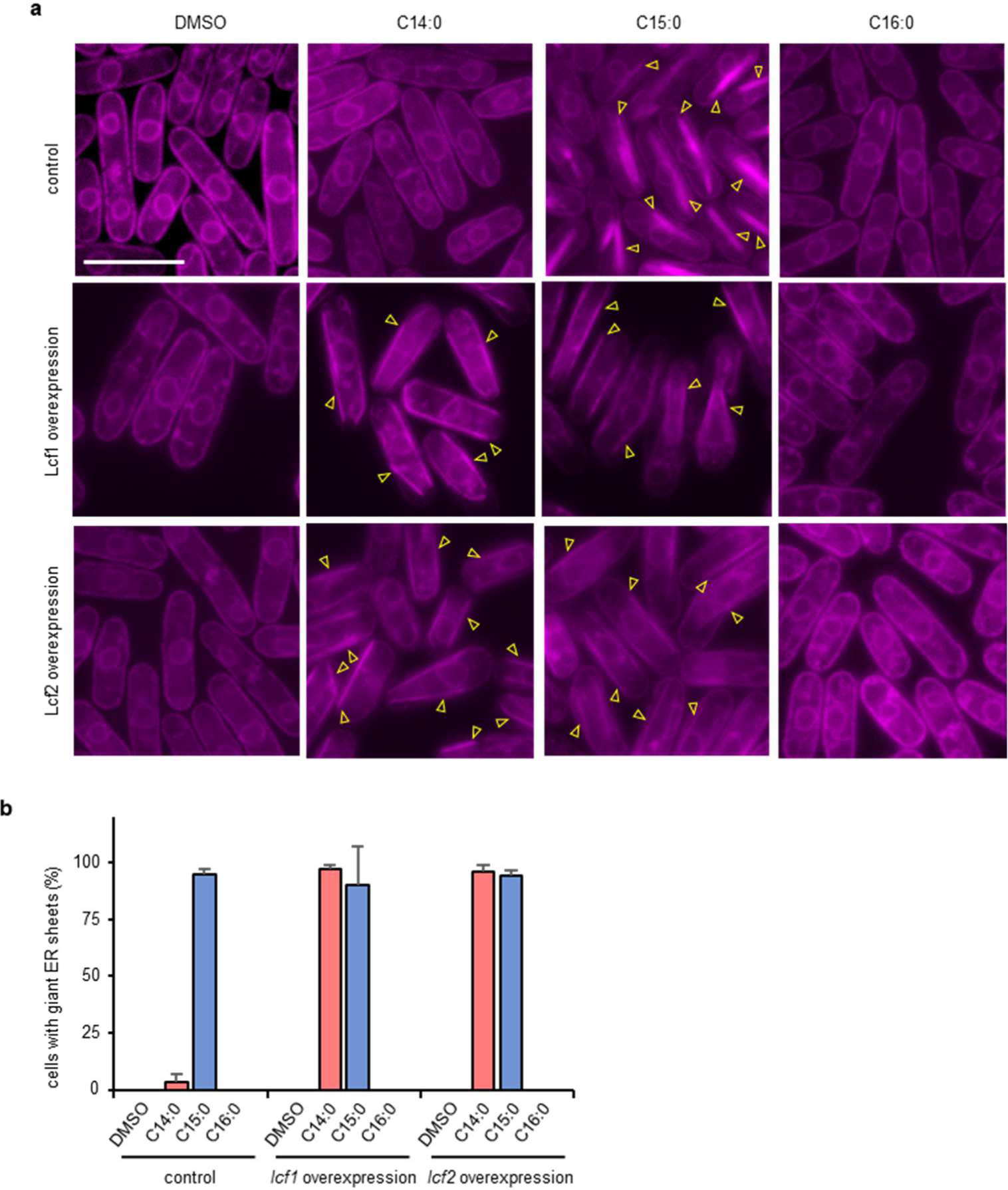
C14:0 induce giant ER sheet formation in cells overexpressing Lcf1 and Lcf2. **a,** The ER in cells overexpressing *lcf1-YFH* and *lcf2-YFH.* Scale bar = 10 µm. **b,** The rate of cells with the giant ER sheets. n > 300 cells from three independent experiments. Error bar = SD.

**Extended Data Figure S18.**
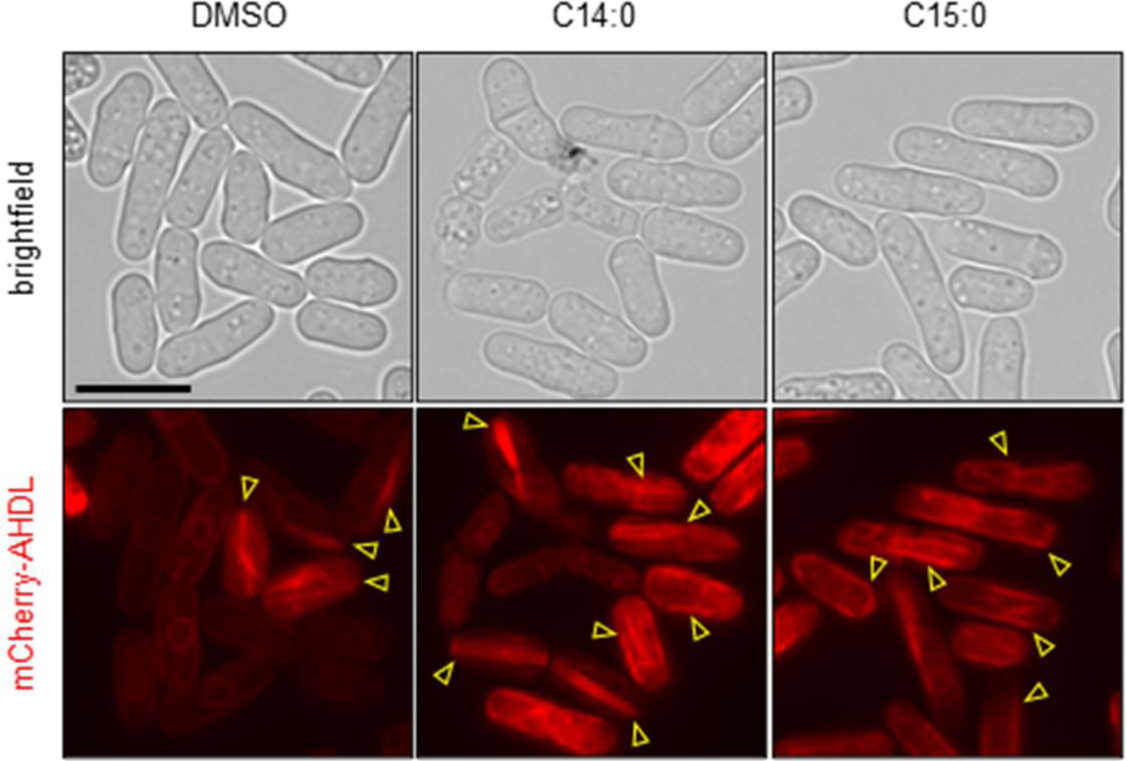
The ER morphology of P *nmt41-ole1* cells. The *ole1* mutant cells expressing mCherry-AHDL were cultivated in MM without or with long-chain FAs and observed under fluorescence microscopy. Yellow arrow heads indicate giant ER sheets.

**Extended Data Table S3.**
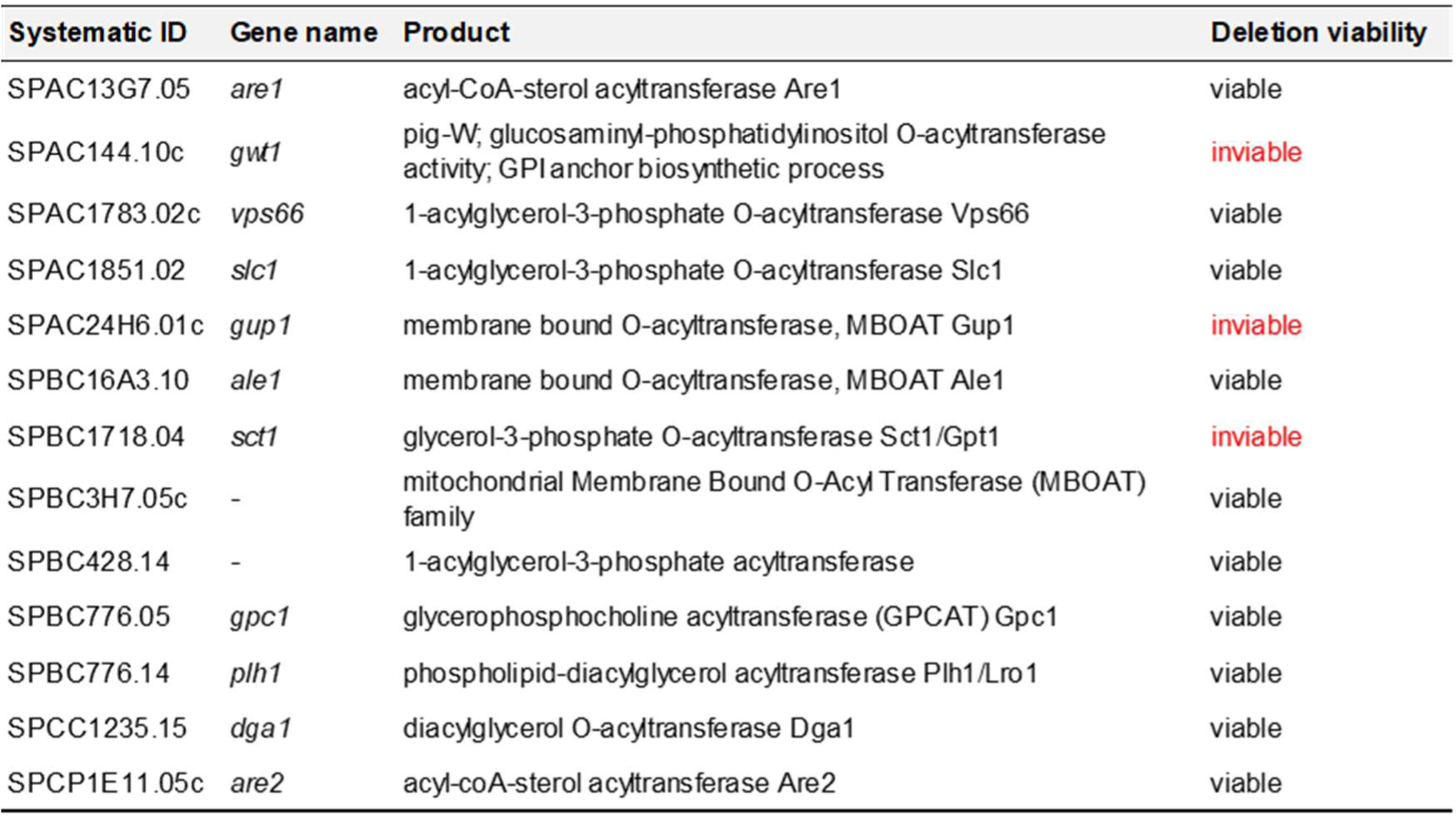
Genes encoding O-acyltransferase in *S. pombe.* Note that acetyltransferases are excluded from the table. Gene names (ID and common name), products, and deletion viability are shown.

| Systematic ID | Gene name | Product | Deletion viability |
| --- | --- | --- | --- |
| SPAC13G7.05 | *are1* | acyl-CoA-sterol acyltransferase Are1 | viable |
| SPAC144.10c | *gwt1* | pig-W; glucosaminyl-phosphatidylinositol O-acyltransferase activity; GPI anchor biosynthetic process | inviable |
| SPAC1783.02c | *vps66* | 1-acylglycerol-3-phosphate O-acyltransferase Vps66 | viable |
| SPAC1851.02 | *slc1* | 1-acylglycerol-3-phosphate O-acyltransferase Slc1 | viable |
| SPAC24H6.01c | *gup1* | membrane bound O-acyltransferase, MBOAT Gup1 | inviable |
| SPBC16A3.10 | *ale1* | membrane bound O-acyltransferase, MBOAT Ale1 | viable |
| SPBC1718.04 | *sct1* | glycerol-3-phosphate O-acyltransferase Sct1/Gpt1 | inviable |
| SPBC3H7.05c | - | mitochondrial Membrane Bound O-Acyl Transferase (MBOAT) family | viable |
| SPBC428.14 | - | 1-acylglycerol-3-phosphate acyltransferase | viable |
| SPBC776.05 | *gpc1* | glycerophosphocholine acyltransferase (GPCAT) Gpc1 | viable |
| SPBC776.14 | *plh1* | phospholipid-diacylglycerol acyltransferase Plh1/Lro1 | viable |
| SPCC1235.15 | *dga1* | diacylglycerol O-acyltransferase Dga1 | viable |
| SPCP1E11.05c | *are2* | acyl-coA-sterol acyltransferase Are2 | viable |

## Notes

### Competing Interest Statement

The authors have declared no competing interest.

